# Killer meiotic driver prevents chromosome loss and enhances fitness in a fungal pathogen

**DOI:** 10.64898/2026.10.04.756499

**Authors:** Gema Puebla Planas, Andrea Doddi, Dilay H. Ayhan, Pilar Gutiérrez Escribano, Li-Jun Ma, Manuel Sánchez López-Berges, Antonio Di Pietro

## Abstract

Fungal pathogens often carry dispensable chromosomes (DCs) that confer a selective advantage in adverse environments or during host infection but can be spontaneously lost under non-selective conditions. How DCs are maintained in evolving populations remains poorly understood. Here we show that a meiotic driver known as Spore Killer (*spok*), located on a giant *Starship* transposon, limits spontaneous loss of a conserved DC in the cross-kingdom pathogen *Fusarium oxysporum*. Transfer of a functional *spok* gene, but not of a killing-deficient mutant allele, to a mitotically unstable DC significantly reduced its loss rate, while *spok* deletion from a stably maintained DC increased loss frequency. Competition experiments between isogenic strains carrying a functional or a killing-deficient *spok* allele revealed a dramatic fitness gain conferred by Spok killing activity, which required transmission by hyphal fusion. Thus, *spoks* and their *Starship* carriers benefit fungal hosts by preventing DC loss and preserving long-term metabolic and pathogenic plasticity.

## Introduction

At least 15% of eukaryotes, including fungi, animals, and plants, possess functionally dispensable but stably propagated supernumerary (“B”) chromosomes^1^. While dispensable chromosomes (DCs) are non-essential for life, they can be beneficial for the organism. In plant pathogenic fungi, DCs are often part of the accessory genome^2^ and confer fitness advantages such as host-specific pathogenicity, resistance to xenobiotics or use of non-preferred carbon sources^3,4^. Some DCs can be horizontally transferred, allowing the recipient strain to cause disease on a new plant species^5,6^. However, because DCs are non-essential their maintenance imposes a fitness cost under non-selective conditions, favouring spontaneous loss^7^. A long-standing question in fungal biology is how DCs are maintained in populations evolving during extended periods in permissive environments, a conundrum analogous to the "plasmid paradox" known in bacteria^8^.

Here we addressed this question in *Fusarium oxysporum*, a fungal pathogen that provokes vascular wilt disease on more than 150 different crop species^9^. Beyond its devastating agricultural impact, *F. oxysporum* can cross the boundaries of host kingdoms, infecting diverse organisms including mammals^10^ and causing life-threatening fusariosis in immunocompromised humans^11^. The genome of *F. oxysporum* is compartmentalized into conserved core and lineage-specific accessory chromosomes, which carry genes involved in host-specific pathogenicity and are rich in Transposable Elements (TEs)^5,6,12^. Most accessory chromosomes, and even some core chromosomes of *F. oxysporum* are conditionally dispensable and can be lost in axenic culture, albeit at very low frequencies^3,4^.

The long-term persistence of DCs in evolving populations suggests the presence of non-Mendelian mechanisms of inheritance^2^. Killer meiotic drivers are selfish genetic elements that employ a poison-antidote system to bias their own transmission by selectively destroying the progeny or neighbouring cells lacking the drive locus^13^. Examples of meiotic killers in fungi include the *wtf* genes from *Schizosaccharomyces pombe*^14^ and the Spore Killer (*spok*) genes in *Podospora anserina*^15^. A recent survey detected an accumulation of *spok* gene homologs in accessory regions of different *Fusarium* lineages, some of which lack a known sexual cycle, leading to the hypothesis that *spoks* could reduce the loss of DCs during vegetative growth^15,16^.

Here we set out to experimentally test the role of a *spok* meiotic driver in DC maintenance and fitness of *F. oxysporum*. By comparing two natural isolates exhibiting dramatic differences in spontaneous loss rate of a conserved DC, we found that the killing function of a *spok* gene, located on a giant *Starship* transposon, significantly enhances DC mitotic stability. Our results establish a key role of these selfish genetic elements in DC maintenance of a clonally propagating fungal pathogen.

## Results

### Two *F. oxysporum* isolates di<er dramatically in mitotic stability of a dispensable core chromosome

Previous experimental evolution studies showed that the clonally evolving *F. oxysporum* f. sp. *lycopersici* (Fol) isolate Fol4287 adapts rapidly to different environments^17^. Here we used an analogous Evolve and Resequence approach to follow DC dynamics in two different *F. oxysporum* strains: the tomato pathogenic isolate Fol4287^18^ and the human cornea isolate MRL8996^19^, that carry distinct DCs^12,20^. Both strains were submitted to ten serial passages on complete (YPDA) or minimal medium plates (MMA). Subsequently, Illumina short read genome sequencing of the evolved populations was performed, and the reads were aligned to the genome assembly of each corresponding wild type strain (Fig. 1). Strikingly, all independently evolved populations of isolate MRL8996 (five on complete and three on minimal medium) had lost chromosome 10, as shown by the absence of reads aligning to this chromosome (Fig. 1a). A synteny analysis between MRL8996 and Fol4287 revealed that MRL8996 chromosome 10 is a conserved core chromosome homologous to chromosome 12 of Fol4287 (Extended Data Fig 1). Of note, Fol4287 chromosome 12 was previously reported to be conditionally dispensable, undergoing spontaneous loss in axenic culture at very low frequencies^3,4^. Here we failed to detect loss of chromosome 12 in the Fol4287 lines passaged on YPDA (5 lines) or MMA plates (4 lines) (Fig. 1b). We conclude that the two *F. oxysporum* isolates MRL8996 and Fol4287 differ significantly in the mitotic stability of a homologous core DC (hereafter called DC10 and DC12, respectively).

**Figure 1.**
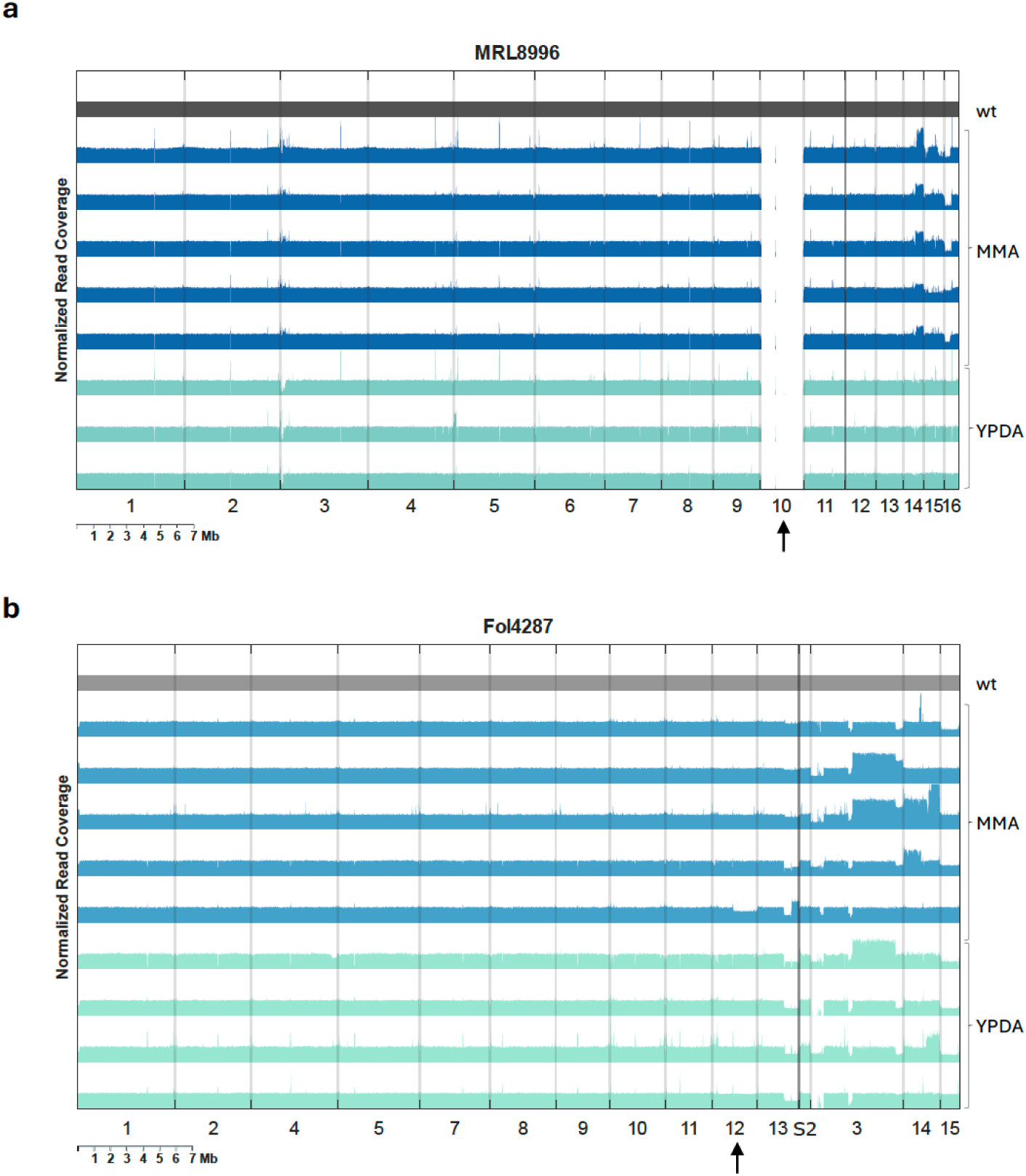
Differential loss of a dispensable core chromosome during experimental evolution of two *F. oxysporum* isolates. **a, b.** Read coverage distribution (10 kb windows) showing loss or duplication of chromosomes or chromosomal regions in multiple independently evolved lines of the *F. oxysporum* strains MRL8996 (**a**) and Fol4287 (**b**). Clonal isolates (wt) were subjected to ten serial passages on minimal (MMA) or complete medium (YPDA) plates. Each passage consisted of spot-inoculating 10^6^ microconidia at the center of the plate and incubating 7 days at 34°C. After passage 10, the evolved lines were subjected to Illumina short read sequencing. Read coverages were normalized first to the median read depth of each sample and then to the wt. Core chromosomes on the left and accessory chromosomes/regions on the right are separated by a black vertical line. Arrows indicate the homologous core chromosomes 10 in MRL8996 and 12 in Fol4287.

### DC10 confers enhanced fitness on non-preferred carbon sources or plant tissue

While DCs are non-essential for survival, they often encode functions that increase fitness in specific environments^3,4^. A gene ontology analysis for categories enriched on MRL8996 DC10 identified functions related to degradation and use of non-preferred carbon sources (hydrolase and xylanase activity, carbohydrate metabolic processes), or with defensive stress responses and xenobiotic degradation (monoxygenase activity, iron ion and heme binding) (Extended Data Fig. 2a). This result is in line with a previous study showing that DC12 contributes to ribose metabolism of Fol4287^3,4^. To test if DC10 facilitates growth of MRL8996 on non-preferred carbon sources, we used RT-qPCR with chromosome-specific primers to monitor its maintenance during five serial passages on MMA plates containing either the preferred carbon source sucrose or the non-preferred sources ribose or polygalacturonic acid. After a single passage on sucrose medium, more than 80% of the population had lost DC10, and after the third passage DC10 was completely lost from the population (Figure 2a). In contrast, populations passaged either on ribose or PGA medium displayed significantly lower loss of DC10, which remained present in more than half of the population after passage 3. This suggests that DC10 contributes to growth of MRL8996 on these non-preferred carbon sources but is dispensable on the preferred carbon source sucrose.

**Figure 2.**
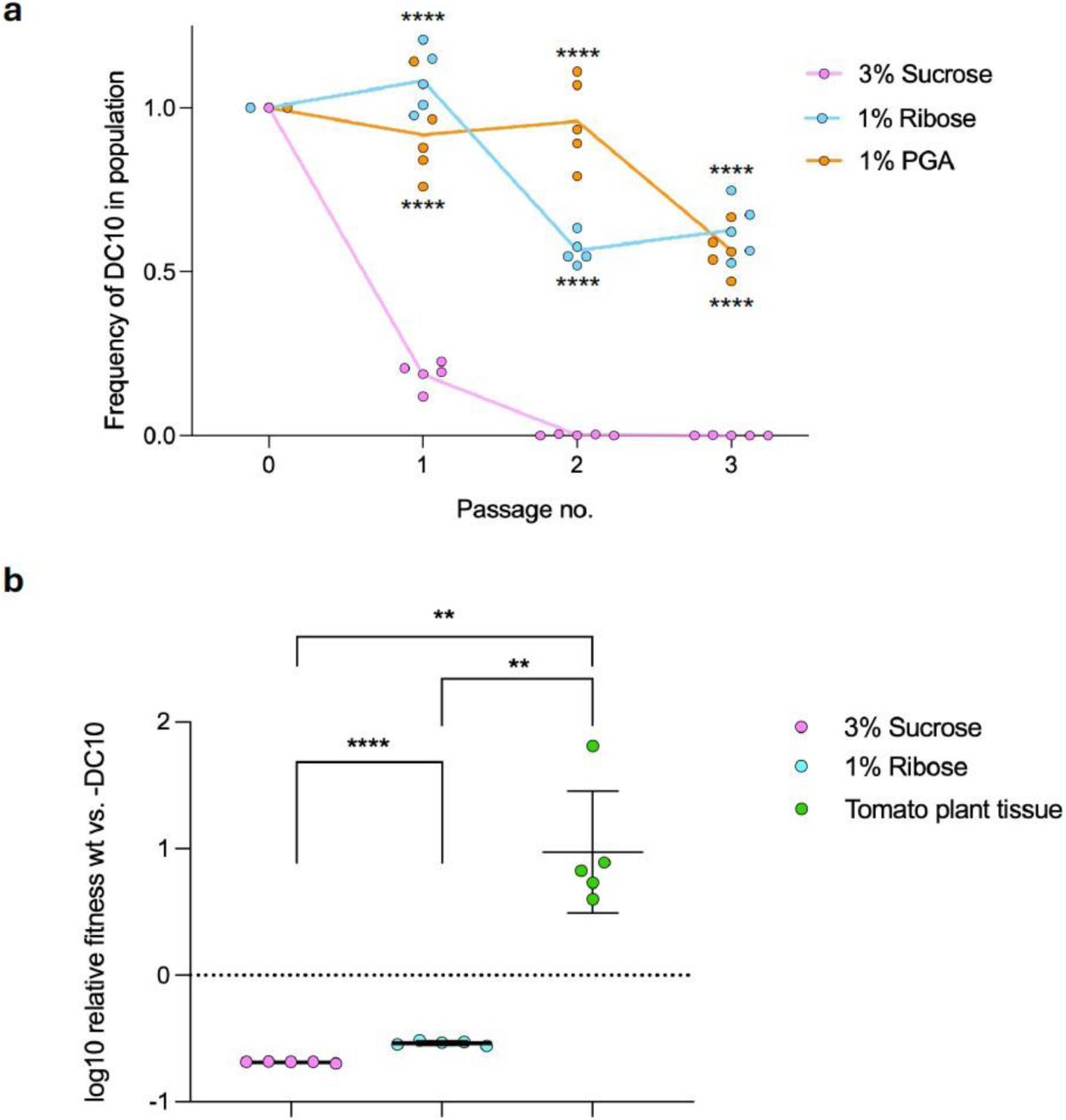
Dispensable chromosome 10 confers a fitness advantage during growth of MRL8996 on non-preferred carbon sources and on plant tissue. **a.** Spontaneous loss of DC10 during growth on different carbon sources. Isolate MRL8996 was submitted to three consecutive passages on plates containing minimal medium (MM) supplemented with the indicated carbon source, by spot-inoculating 10^6^ microconidia at the center of the plate. After 4 days of incubation at 34°C or 7 days of incubation at 28°C for the plant tissue, conidia from the entire colony were collected for DNA extraction and for inoculation of the next passage. For each carbon source, five independent lines were passaged. The frequency of DC10 in the passaged populations was measured by real-time qPCR of total genomic DNA extracted from microconidia, using a specific primer pair, and normalized to the *actin* gene located on the core chromosome 5. ****, *p* < 0.0001 versus sucrose, according to ordinary 2-way ANOVA. Each data point represents an independently passaged line. **b.** DC10 increases fitness on non-preferred, but not on preferred carbon sources. Scatter plots show the log10 relative fitness of the MRL8996 wild type strain versus a strain that spontaneously lost DC10 (-DC10), calculated from competition experiments. A 1:1 mix of microconidia from the two strains was either spot-inoculated at the center of a plate containing MM with sucrose or ribose as the carbon source (10^6^ conidia in total) or distributed on top of a tomato plant previously killed by immersion in liquid N2 (2.5×10^5^ conidia in total). After 4 or 7 days of incubation, respectively, at 28°C, total microconidia from the plate or the total plant biomass were harvested, and the frequency of DC10 in the MRL8996 population was measured as described in **a**. Error bars, SD. ****, *p* < 0.0001, **, *p* < 0.01 according to Brown-Forsythe and Welch ANOVA tests. Each data point represents an independent competition experiment.

To quantify the fitness contribution of DC10 on different carbon sources, we performed pairwise competition experiments between the MRL8996 wild type strain and a monosporic isolate that had spontaneously lost DC10 (-DC10), as confirmed by PCR and Nanopore sequencing (Extended Data Fig. 2b). The relative fitness of the wild type versus -DC10 was significantly higher on ribose than on sucrose medium, and around 9-fold higher (geometric means) during growth on tomato plant tissue (Fig. 2b). These findings indicate that DC10 significantly contributes to fungal fitness of *F. oxysporum* MRL8996 on non-preferred carbon sources, particularly during growth on plant tissue.

### Fol4287 DC12 carries a giant *Starship* transposon harbouring a spore killer gene

To understand the genetic basis underpinning the dramatic difference in mitotic stability of DC12 in Fol4287 versus its homolog DC10 in MRL8996, we performed a sequence alignment between the homologous chromosomes of the two strains. This revealed the presence of a 92.5 kb region in Fol4287 DC12 that is absent in the MRL8996 homolog (Extended Data Fig. 3a). The sequence of the insertion exhibits the hallmarks of giant *Starship* transposons, including a putative tyrosine recombinase known as Captain (Pfam domain DUF 3435) in the first position^21^. Within the *Starship* element, we observed a *spok* gene whose predicted protein product displays the characteristic tripartite domain structure of spore killer meiotic drivers^15,16^. Alignment with the amino acid sequence of Spok3 from *P. anserina* detected significant homology in the nuclease killing and the kinase resistance domains (Extended Data Fig. 3b). Importantly, the lysin residue at position 240, which is essential for Spok killing activity^15,16^, is positionally conserved (Extended Data Fig. 3b, c), suggesting that Fol4287 Spok is likely functional.

Because the *Starship/spok* insertion is located on a dispensable core chromosome of Fol4287, we asked whether it is conserved in the homologous chromosomes of other *Fusarium* strains. Screening of 1,252 publicly available genome assemblies for sequences homologous to Fol4287 chromosome 12 identified alignments in 773 assemblies, 421 of which covered at least 70% of the length of Fol4287 chromosome 12 and were retained for further analysis (Extended Data Fig. 4a). In 226 of these 421 isolates, *Starship* was not detected, in 116 isolates *Starship* was present with < 50% coverage, in 57 isolates with 50-75% coverage, and only in 22 isolates *Starship* was apparently intact, containing both the *captain* and the *spok* gene. Most of these 22 isolates belong to the formae speciales *lycopersici* (tomato pathogen; n=12) and *cubense* (banana pathogen; n=6), forming a tight, near-monophyletic group in an identity-based maximum-likelihood tree of chromosome 12 homologs (Extended Data Fig. 4a). In contrast, the *spok* genes identified outside of an intact *Starship* (n=94) are phylogenetically dispersed.

To ask whether this phylogram is specific for DC12 homologs or reflects a broader genomic relationship, we performed the analogous analysis for another small core chromosome of Fol4287, chromosome 10 (homologous to MRL8996 chromosome 8; see Extended Data Fig. 1). The 468 isolates that passed the 70% identity threshold for Fol4287 chromosome 10 displayed a remarkably similar distribution as in the phylogram of DC12, including the same phylogenetically related group of isolates (Extended Data Fig. 4a). Together, these findings are consistent with a monophyletic origin of the *Starship/spok* insertion on DC12 in a common ancestor of this group of isolates.

### Transfer of a functional *spok* gene to MRL8996 DC10 increases its mitotic stability under non-selective conditions

To experimentally test the role of Spok in DC maintenance, we cloned the *spok* gene from DC12 of Fol4287 and transferred it to a specific locus (*pg1*) on the homologous DC10 of MRL8996 (Extended Data Figs. 5 and 6). As a negative control, we created a mutated *spok* allele in which the conserved K240 residue (see Extended Data Fig. 3b, c) had been replaced with an alanine to abolish killing activity^15,16^. To allow accurate monitoring of the presence or loss of DC10, the hygromycin resistance cassette and three tandem copies of the green fluorescent reporter gene *mclover3* (*3xclover*) were included in the construct (Extended Data Fig. 6). The obtained transformants, named MRL8996::*clover+spok* and MRL8996::*clover+spok^K240A^*, respectively, were submitted to five consecutive passages on minimal medium with sucrose as the carbon source, performing ten independently passaged lines per strain. After passages 1, 3 and 5, the frequency of DC10 in the evolved populations was measured using three different methods: 1) counting of hygromycin-resistant colonies; 2) RT-qPCR of total genomic DNA with DC10-specific primers; and 3) quantifying the proportion of fluorescent cells by flow cytometry. The three independent methods detected a progressive loss of DC10 in the evolved lines carrying the killing-deficient *spok^K240A^* allele, reaching a mean loss rate of 50-70% after passage 5 (Fig. 3a-c). In stark contrast, the lines carrying the wild type *spok* allele inserted on DC10 displayed a significantly reduced loss rate of 10-20%. Importantly, the three distinct approaches of DC quantification provided similar results, although flow cytometry proved to be the most efficient and reliable method due to its superior speed and sampling depth (100,000 events per sample). Overall, these results demonstrate that insertion of a functional *spok* gene is sufficient to significantly increase the mitotic stability of the previously highly unstable DC10 in *F. oxysporum* MRL8996.

**Figure 3.**
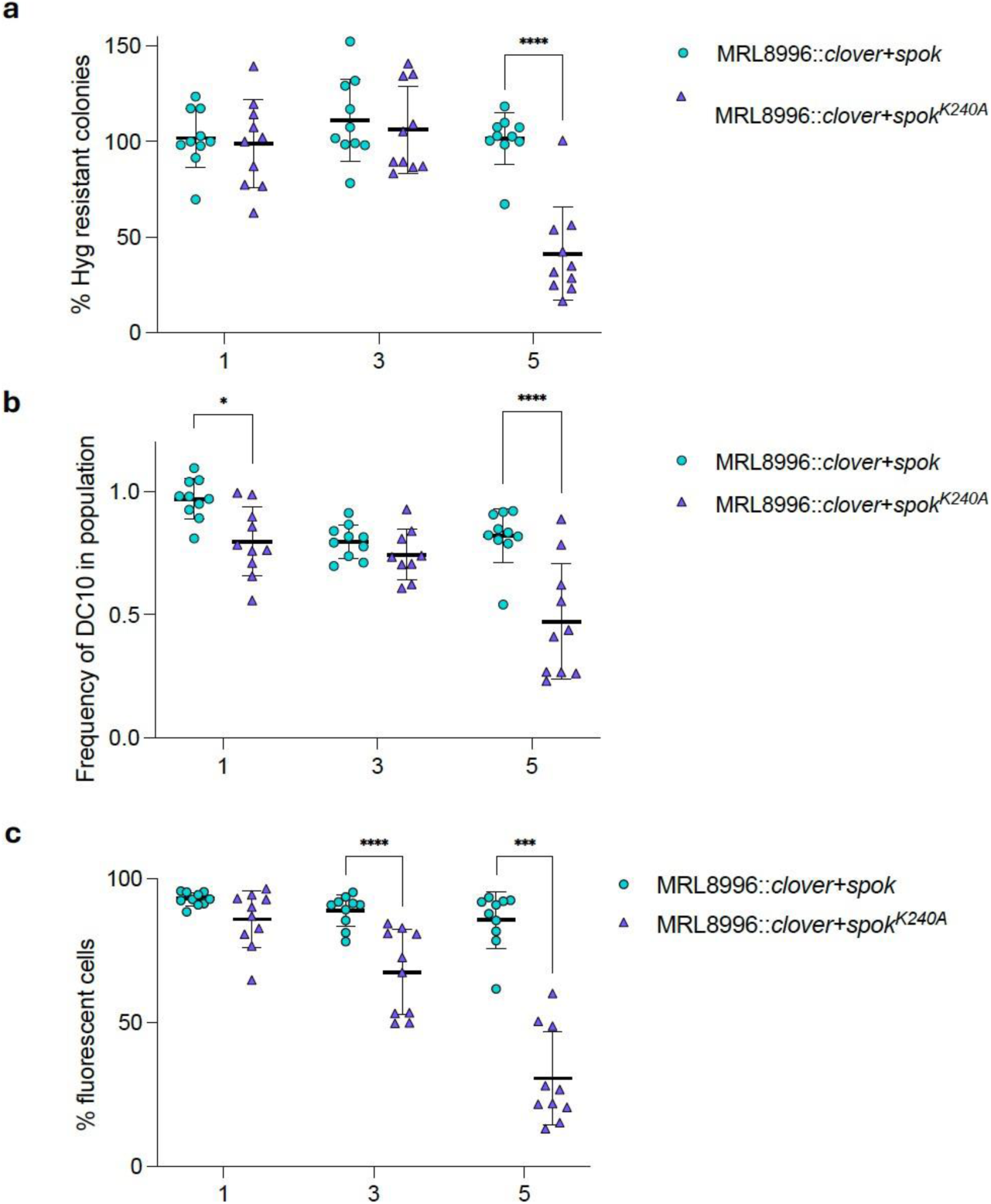
Insertion of a functional *spok* gene reduces spontaneous loss of DC10 in MRL8996. **a-c.** Spontaneous loss of DC10 during plate passages of the indicated MRL8996 strains carrying either a wild type or a killing-deficient *K240A* allele from Fol4287 *spok*, as well as the *hyg^R^* and the green fluorescent *3xclover* marker genes (see Extended Data Fig. 6). Strains were submitted to five consecutive passages at 34°C on minimal medium with sucrose as the carbon source. For each genotype, 10 independent lines were passaged. After passages 1, 3 and 5, the frequency of DC10 in the population was measured by (**a**) counting colony forming units (CFUs) on PDA without or with hygromycin (2 replica plates per sample); (**b**) RT-qPCR of total genomic DNA extracted from microconidia using specific primers and normalized to the actin gene (3 technical replicates); or (**c**) fluorescence flow cytometry (100’000 microconidia per sample). Error bars, SD. ****, *p* < 0.0001, **, *p* < 0.01, *, *p* < 0.05 according to ordinary 2-way ANOVA. Each data point represents an independently passaged line.

### Spok killing activity counteracts spontaneous DC loss

The progressive decline of DC10 frequency in plate-passaged populations of MRL8996 (see Figs. 2 and 3) likely reflects two mechanistically distinct processes: 1) mitotic defects resulting in cells that have lost the DC (spontaneous loss rate); and 2) competition between the cells of the population that carry or lack the DC, leading to shifts in DC frequency (fitness/selection effect). Spok is predicted to contribute to the second process by selectively killing the cells that have lost the DC harbouring *spok*, thereby favouring selection for DC maintenance. However, the killing effect of Spok could be partially masked by environmental parameters such as the presence of a preferred carbon source, which selects against DC presence (see Fig. 2). To focus on the Spok killing effect while minimizing other factors, we compared the DC10 loss rates between the MRL8996::*clover+spok* and MRL8996::*clover+spok^K240A^* strains after one single plate passage. Flow cytometry analysis of ten independent lines per strain detected a significantly lower mean loss rate in MRL8996::*clover* carrying a wild type *spok* gene compared to the isogenic strain carrying the killing-deficient *spok^K240A^*allele (Fig. 4a). This result indicates that Spok killing function successfully counteracts spontaneous DC10 loss.

**Figure 4.**
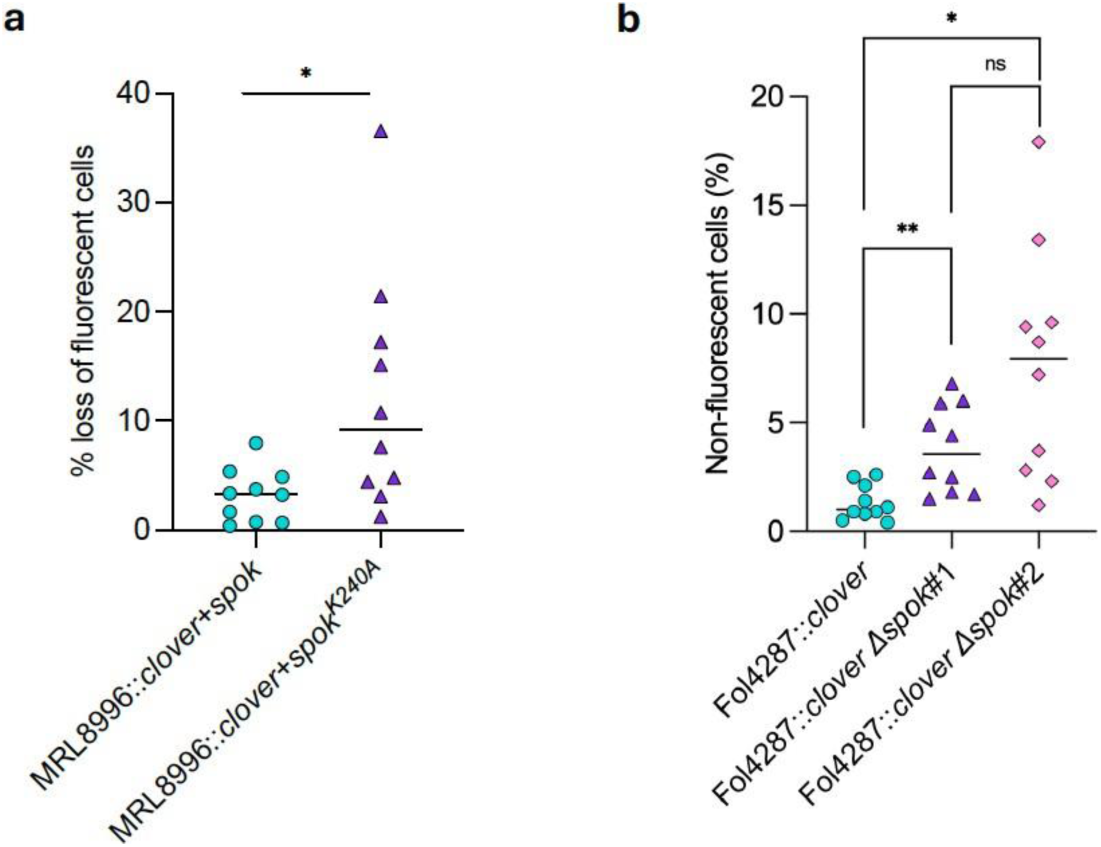
Loss of Spok-mediated killing leads to increased DC loss rate. Spontaneous loss rate of DC10/DC12 in the indicated strains after a single passage at 34°C on minimal medium plates with sucrose as the carbon source was quantified by fluorescence flow cytometry (100’000 microconidia per sample). Note that all strains carry the *hyg^R^* and green fluorescent *clover* marker genes inserted in the dispensable chromosome (see Extended Data Figs. 5 and 6) **a.** Loss rate of DC10 in transformants of MRL8996 carrying either a wild type or a killing-deficient *K240A* allele of the Fol4287 *spok* gene. **b.** Loss rate of DC12 in the Fol4287 wild type strain and in two independent Δ*spok* deletion mutants. **, *p* < 0.01, *, *p* < 0.05 according to Brown-Forsythe and Welch ANOVA tests. Each data point represents an independent plate passaged line.

To independently confirm the role of Spok in DC maintenance, we used a complementary approach consisting of deleting the *spok* gene from Fol4287 DC12. Targeted gene knockout was performed in a Fol4287::*clover* background, carrying the *3xclover* fluorescent marker and the hygromycin resistance cassette inserted at the *pg1* locus on DC12 (Extended Data Figures 7 and 8). Two independent Fol4287::*clover* Δ*spok* mutants and the Fol4287::*clover* wild type *spok* strain were subjected to a single plate passage, and the presence of DC12 in the population was quantified by flow cytometry. Analysis of ten independent lines per strain revealed a significantly higher loss frequency of DC12 in the two Δ*spok* mutants compared to the wild type *spok* strain (Fig. 4b). Together, these results demonstrate that Spok killing activity significantly reduces short-term DC loss rate in two different *F. oxysporum* strains.

### Spok-mediated killing confers increased competitive fitness mediated by hyphal fusion

Meiotic drive spore killers confer a selective advantage to the host organism by increasing their own frequency in the population, although the destruction of non-killer sibling spores can impose a fitness cost^22^. To date, the fitness effect of spore killers in clonally evolving fungi such as *F. oxysporum* remains unexplored. The availability of near-isogenic strains carrying either a functional or a killing-deficient *spok* allele allowed us to address this question through pairwise competition experiments between fluorescent and non-fluorescent strains. We first competed MRL8996::*clover+spok* or MRL8996::*clover+spok^K240A^* against the non-fluorescent MRL8996 wild type or -DC10 strain. Competition assays were carried out on MMA plates with either sucrose or ribose as carbon source, or on PDA. The proportion of fluorescent versus non-fluorescent cells at the start and the end of the experiment was measured by flow cytometry and used to calculate the effect of Spok presence/absence on competitive fitness. In all experimental conditions tested, the relative fitness of the MRL8996::*clover+spok* strain was at least an order of magnitude higher compared to the MRL8996::*clover+spok^K240A^* strain, independent of the carbon source or of the presence/absence of DC10 in the competitor strain (Fig. 5a). Similarly, the Fol4287::*clover* strain showed dramatically increased fitness compared to the two Fol4287::*clover* Δ*spok* mutants, when competed against the non-fluorescent Fol4287 wild type (Fig. 5b). Importantly, colony growth speed of these strains did not differ significantly, thereby ruling out a direct effect of Spok on hyphal growth (Extended Data Fig. 9). Together, these results establish that Spok confers a dramatic fitness advantage during vegetative growth of *F. oxysporum*, likely by selectively killing the cells of the colony that lack a functional *spok* gene.

**Figure 5.**
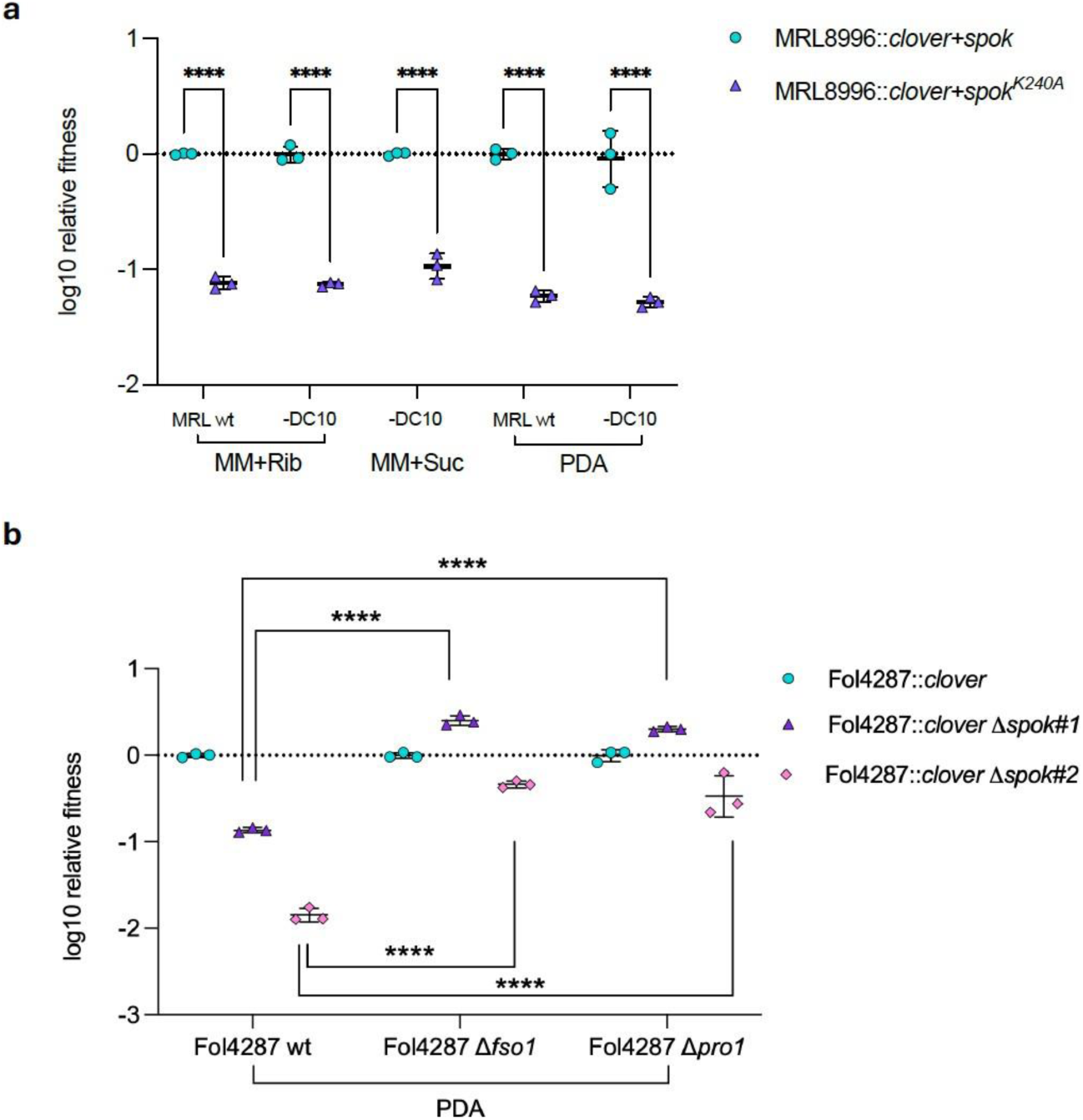
Spok-mediated killing confers increased fitness and requires hyphal fusion. Scatter plots showing the log10 relative fitness of the indicated fluorescent strains, as determined from competition experiments against the non-fluorescent strains shown in the x-axis. **a.** MRL8996::*clover* strains carrying either a wild type or a killing-deficient allele of the Fol4287 *spok* gene inserted in DC10 were spot-co-inoculated (1:1 mix) together with the non-fluorescent MRL8996 wild type strain (MRL wt) or with a derivative thereof lacking DC10 (-DC10) on plates containing either PDA or MM supplemented with 1% ribose or 3% sucrose as the carbon source. **b.** The Fol4287::*clover* strain or two independent Fol4287::*clover* Δ*spok* mutants were spot-coinoculated (1:1) together with the non-fluorescent Fol4287 wild type strain or with hyphal fusion-deficient mutants lacking either the *fso1* or the *pro1* gene on PDA plates. After 3 days incubation at 28°C, total microconidia from the colonies were harvested and the proportion of fluorescent versus non-fluorescent cells at the start and the end of the experiment was determined by flow cytometry (n=100’000 conidia per sample in (**a**), (60’000 in (**b**)), used to calculate the relative fitness of the fluorescent strains, normalized against MRL::*clover+spok* (**a**) or Fol4287::*clover* (**b**) (see Methods for details on fitness calculations). Error bars, SD. ****, *p* < 0.0001 according to Dunnett two-way ANOVA. Each data point represents an independent plate competition experiment.

A key question is how Spok killing is enacted within a mixed colony of vegetatively growing near-isogenic strains. A potential mechanism is vegetative hyphal fusion, a process that physically connects individual mycelial networks, allowing efficient exchange of cytoplasm and nutrients, but also of selfish genetic elements^23^. To ask whether hyphal fusion is required for Spok-mediated killing and for the resulting fitness gain, we competed the Fol4287::*clover* strain or the Fol4287::*clover* Δ*spok* mutants against two isogenic deletion mutants lacking the *fso1* or the *pro1* gene, either of which is required for efficient hyphal fusion of Fol4287^24,25^. Strikingly, the dramatic fitness defect of the Δ*spok* strains observed during competition with the fusion-competent wild type strain was abrogated when they were competed against the fusion-deficient mutants (Fig. 5b). Together, these findings demonstrate that Spok-mediated killing confers a strong fitness advantage during vegetative hyphal growth of *F. oxysporum*, which is mediated by hyphal fusion.

## Discussion

The remarkable stability of DCs in fungal populations has long represented an evolutionary puzzle. Our findings provide a mechanistic explanation for this conundrum, by demonstrating that a spore killer meiotic driver encoded on a dispensable core chromosome efficiently reduces its spontaneous loss. Spok acts as a stabilizing force by selectively eliminating the cells that have lost the DC, thereby contributing to its maintenance and preserving long-term metabolic plasticity of *F. oxysporum*. Our study experimentally demonstrates the functional impact of Spok via two independent genetic approaches. First, introduction of the *spok* gene from Fol4287 DC12 to the homologous, inherently unstable DC10 of MRL8996 drastically reduced its spontaneous loss rate, demonstrating that this spore killer element is both functional and transferable between strains. Second, targeted deletion of *spok* from the stably maintained DC12 in Fol4287 significantly increased DC loss rate. The ability to quantify chromosome loss after a single passage was essential for decoupling loss rate from subsequent environmental selection acting on the evolving population. In our hands, flow cytometry with strains expressing a highly fluorescent *3xclover* marker was the most precise and statistically robust method to quantify DC loss rate in single culture, as well as fitness differentials in competition experiments.

Spok killing activity was essential for the DC-stabilizing effect, which was abolished by a single point mutation in the conserved Spok killing domain. Given the widespread distribution of *spok* genes in *Fusarium* and other Pezizomycotina^15,16^, we speculate that their role in DC maintenance could be broadly conserved across these fungal lineages. Intriguingly, the killer-antidote protective mechanism of fungal Spoks mirrors the evolutionarily ancient toxin–antitoxin system encoded on bacterial plasmids, which maintains plasmid stability through post-segregational killing of plasmid-free cells^26^, contributing to maintenance and dissemination of antibiotic resistance^27^.

Fungal DCs often confer context-dependent fitness advantages. A classic example is a small DC of *Fusarium solani* harbouring the pisatin demethylase gene whose product detoxifies the antimicrobial compound pisatin produced by garden pea, thereby allowing the fungus to specifically colonize pea plants^3,4^. Here we found that DC10 dramatically increases the competitiveness of *F. oxysporum* MRL8996 during growth on dead plant tissue. The fact that this cornea isolate, which appears to be adapted to clinical settings^28^, maintains a DC enriched in genes involved in mobilization and use of non-preferred carbon sources and in xenobiotic degradation, suggests that it retains the capacity for a saprophytic lifestyle based on colonization of plant biomass, allowing it to thrive in environmental or agricultural niches. On the other hand, a MRL8996 strain that had lost DC10 displayed increased fitness on the preferred carbon source sucrose, strongly suggesting that maintenance of DC10 entails significant metabolic costs and explaining its progressive loss observed under non-selective conditions. Together, these findings offer fascinating insights into DC dynamics and fitness trade-offs in *F. oxysporum* during adaptation to different ecological niches.

Intriguingly, the *spok* gene on Fol4287 DC12 is located within a giant *Starship*TE, a constellation first reported in the *Starship Enterprise* from *P. anserina*^29^ and subsequently found in a wide variety of fungi, including *Fusarium*^16,21^. In a recent survey of *spok* genes in *Fusarium* genomes, 33 (8%) of the 408 *spoks* analyzed were located within *Starships*^16^. The cooperation between a killer meiotic driver and a giant transposon represents a powerful alliance of two selfish genetic elements that is mutually beneficial, setting the basis for a long-term parasitic alliance^30^. Intriguingly, our results demonstrate that the *Starship/spok* alliance also provides at least two important benefits to the host organism: 1) contributing to the evolutionary maintenance of DCs, and 2) dramatically enhancing the organism’s overall fitness during physical interaction with compatible fungal strains.

DC12, which harbours the *Starship/spok* insertion in Fol4287, is a dispensable fast core chromosome^3,4^. This term designates the three smallest core chromosomes of *F. oxysporum,* which have some hallmarks reminiscent of accessory chromosomes such as a higher sequence divergence and an enrichment of histone H3 lysine 27 trimethylation (H3K27me3) associated with facultative heterochromatin^31^. Our analysis of 421 *Fusarium* genomes in which DC12 is conserved revealed that most isolates carrying an intact *Starship* with a *spok* gene clustered within a single clade, strongly suggesting a monophyletic origin of the *Starship/spok* insertion event, which likely occurred in a common ancestor. This contrasts with the findings reported for the *spok* genes found on accessory chromosomes, which suggested a polyphyletic origin mediated by horizontal transfer^16^ .A recent analysis detected a significant enrichment of *spok* genes on accessory regions, yet the *spoks* found inside of a *Starship* were mostly located on core chromosomes (including the *spok* gene *FOXG_14774* on Fol4287 DC12) or on unknown genomic compartments, with most of them falling within two defined clusters^16^. Together, these findings indicate that the *Starship/spok* module in *F. oxysporum* is predominantly located on homologs of Fol4287 fast core DC12.

A striking result from competition assays between fluorescent and non-fluorescent strains was the strong competitive advantage conferred by the presence of a single *spok* gene. The capacity of Spok to increase fungal fitness required both its killing activity as well as a process known as vegetative hyphal fusion. In environments where different *F. oxysporum* strains compete for limited space and nutrients, Spok-mediated killing could thus provide a crucial ecological advantage by effectively eliminating the nuclei of rival lineages upon hyphal fusion, thereby securing genetic dominance within the heterokaryon. In addition, this strategy would enable resource plundering, similar to nuclear parasitism by *Neurospora crassa* cheater strains^32,33^. A potential limitation of this Spok-mediated fitness gain is heterokaryon incompatibility, a defence system that protects fungi against unauthorized somatic fusion via non-self recognition of different *het* loci, triggering programmed cell death^34^. However, previous studies suggest that the incompatibility response is suppressed under certain environmental or developmental conditions, including starvation^35^ or early stages of colony formation consisting of conidial anastomosis tubes (CATs)^36^. While CAT fusion has been reported in Fol4287^24^, it remains to be investigated whether this mechanism expands the range of Spok-mediated fitness gain across distantly related fungal lineages. On the other hand, similar to *spoks*, the *het* genes that trigger heterokaryon incompatibility can be captured and mobilized by *Starships*^21,37^, raising the intriguing possibility that *Starships* simultaneously carrying *spok* and *het* genes could confer a dual ability to the host for outcompeting rival lineages through *spok*-mediated nuclear killing, while protecting itself from invasion by potential competitors and genetic parasites.

## Methods

### Strains and culture conditions

The tomato pathogenic isolate *Fusarium oxysporum* f. sp. *lycopersici* 4287 (Fol4287)^18^ and the human cornea isolate *Fusarium oxysporum* MRL8996 (NRRL 47514)^19^ were used throughout the study. Fungal strains were routinely stored as microconidial suspensions in 30% glycerol (v/v) at - 80°C. For microconidia production and DNA extraction, strains inoculated from -80°C stocks were grown for 3 days in Potato Dextrose Broth (PDB) at 28°C and 170 rpm and collected by filtration using Monodur filters (pore size, 10-15 μm). When needed, hygromycin (20 μg/mL) or nourseothricin (2.5 μg/ml) were added.

For determination of growth rate, 5×10^4^ freshly obtained microconidia were spot-inoculated at the center of a Potato Dextrose Agar (PDA) plate and incubated at 28°C. Plates were imaged on days 3 and 7 after inoculation. The area of the fungal colony was measured using the MultiGauge software (FujiFilm Corporation, Valhalla, NY, USA) and growth rate was calculated as the net increase in colony area between days 3 and 7. Three replicate plates were performed for each fungal strain and condition.

Serial passaging experiments were performed as described^17^. Briefly, 10^6^ freshly obtained microconidia were spot-inoculated at the center of Yeast Peptone Dextrose Agar (PDA) or Minimal Medium Agar (MMA) plates. Standard MMA contained 3% (w/v) sucrose as the sole carbon source. For some serial passaging experiments, the non-preferred carbon sources 1% (w/v) D-ribose or 1% (w/v) polygalacturonic acid (pga) were used. pH of the plates was adjusted to 7.4. After 7 days of incubation at 34°C, the microconidia from the entire colony were collected by adding 2 ml of sterile distilled water, scraping with a sterile spatula and filtered into a tube with sterile cotton. The concentration was determined using a TC20 Automated Cell Counter (Bio-Rad, Madrid, Spain), adjusted to 10^8^ spores/ml, and 10 µl of this suspension were used to spot-inoculate a fresh plate for the subsequent passage. After each passage, microconidia were stored at -80°C for subsequent analysis.

### Nucleic acid manipxulations

Genomic DNA was extracted from fungal mycelia or microconidia using the CTAB method^38^. The quantity and quality of DNA were determined using a NanoDrop ND-1000 spectrophotometer (NanoDrop Technologies, Wilmington, DE, USA) at 260 nm and 280 nm, and by gel electrophoresis in 0.7% (w/v) agarose gels.

PCR reactions were performed using Expand High Fidelity PCR System (Roche, Barcelona, Spain) for <5 kb fragments, or iProof High Fidelity DNA Polymerase (Bio-Rad) for >5 kb fragments. BioTaq DNA Polymerase (Meridian Bioscience, Newtown, Ohio, USA) was used for routine PCRs and generation of Southern blot probes. All PCR reactions were performed in a T100 Thermal Cycler (Bio-Rad). For real time (RT) quantitative PCR (qPCR), the FastStart Essential DNA Green Master kit (Roche) and a CFX Connect Real-Time System (Bio-Rad) were used according to the manufactureŕs instructions.

### Generation of DNA constructs

The isolates Fol4287 or MRL8996 were used as genetic backgrounds for all fungal strains generated in the study (Supplementary Table S1). Plasmids and PCR primers used are listed in Supplementary Tables S2 and S3, respectively. The different genetic constructs targeting the *pg1* locus (*FOXG_14695*)^39^ were assembled through homologous recombination in yeast^40^. Briefly, PCR was used to obtain the individual fragments as follows: for the Fol4287*::clover* strain: the green fluorescent marker *3xclover* (amplified from pUC57-Fo-mclover3^41^ using primers gpda15N and SV40 R), the hygromycin resistance cassette (amplified from Pan-7.1^42^ using primers gpda15N and HygRevClover) and the 5’- and 3’-flanking regions of the *pg1* gene (amplified from genomic DNA of Fol4287 using primer combinations pg1-15T/pg1-10FH and pg1-7F/pg1-13T, respectively); for the MRL8996*::clover+spok* and MRL8996*::clover+spok^K240A^* strains, the same primer combination was used to amplify the 3x*clover* cassette and the hygromycin resistance cassette. In addition, the *spok* sequence was amplified from genomic DNA of Fol4287 using the primers Spok 5’For/ Spok 3’ Rev Hyg. For the 5’ and 3’ flanking regions of the *pg1* gene (amplified from genomic DNA of MRL8996), the primers used were pg1-15T/pg1-10-MS and pg1-7M/pg1-13T, respectively. All fragments carried at least 30 bp of overlapping sequence. For yeast assembly, the plasmid noR_RFP-T^43^ linearized with *Hind*III was used as the backbone. After assembly in yeast, plasmid DNA was extracted, transformed into *Escherichia coli,* extracted using the CTAB protocol^44^, and treated with the restriction enzyme *Pme*I to release the linear construct of interest used for protoplast transformation. Schematic representations of each construct are shown in Extended Data Figures 6 and 7.

To generate the MRL8996*::spok^K240A^* mutant allele, a point mutation was introduced into the wild-type *spok* gene (*FOXG_14774*) using inverse PCR^45^. Schematic representation of the construct is shown in Extended Data Figure 8. Briefly, the Fol4287 *spok* gene was amplified using the primer combination Spok 5’ For/Spok 3’ Rev and cloned into the pSpark-I vector (Canvax Biotech, Valladolid, Spain). The overlapping primers MutSpokFor and MutSpokRev were then used to introduce the K240A mutation, previously shown to eliminate killing function in the *Podospora anserina spok3* gene^15^. The mutated *spok* allele was confirmed by Sanger sequencing, using the oligonucleotide SpokForSeq located 325 bp upstream of the point mutation.

Targeted deletion of the *spok* gene in the Fol4287*::clover* background was done by homologous gene replacement as described^46^. The nourseothricin resistance cassette, amplified from pAN-8.1^47^ with primers M13For/M13Rev, was fused to the 5’- and 3’-flanking sequences of the *spok* gene, amplified with primer combinations Spok 5’ For n/Spok 5’ Rev Nat and Spok 3’ For Nat/Spok 3’ Rev n, respectively.

### Fungal transformation and analysis of transformants

Targeted integration of the DNA constructs as well as gene deletions were performed by protoplast transformation^48^. Transformants were purified through two rounds of monoconidial isolation in the presence of the selective agent^39^, and the correctness of the integration was confirmed by Southern blot analysis^46^, using probes for *3xclover* (primers CLFor/3xFlagRev).

A MRL8996 strain that had lost DC10 (MRL8996 -DC10) was isolated from plate-passaged line of the MRL8996::*spok^K240A^* strain. The absence of DC10 was confirmed by Nanopore sequencing (Extended Data Figure 2b). A hygromycin resistant derivative thereof (MRL8996 *-*DC10*::hyg*) was obtained by transforming protoplasts with the hygromycin resistance cassette.

### Quantification of DCs loss in fungal strains

Three distinct approaches were used to quantify the loss of DC12 in Fol4287, or of its homolog DC10 in MRL8996: 1) counting of hygromycin-resistant colonies; 2) RT-qPCR analysis; and 3) fluorescence flow cytometry. To count hygromycin-resistant colonies, 50 μl aliquots containing approximately 100 microconidia were uniformly spread on PDA plates with or without hygromycin, using sterile glass beads. After 3 days of incubation at 28°C, the number of colonies in the absence or presence of hygromycin was determined, and the DC loss rate was calculated as the ratio between the total number of hygromycin-resistant colonies and the total number of colonies on non-selective plates. Two technical replicates per strain were performed. For qPCR analysis, genomic DNA was extracted from freeze-dried microconidia using the CTAB method indicated above. During DNA extraction from inoculated tomato material, plant polyphenols were removed as described^49^ with some modifications. Briefly, samples were mixed 1:1 with AMPure XP beads (Beckman Coulter), incubated for 10 minutes at room temperature (RT), and placed on a magnetic rack (Thermo Fisher) until the beads were completely retained. After discarding the supernatant, the beads were washed three times with 200 μl 70% (v/v) EtOH, resuspended in 25 μl distilled water, and incubated for 10 minutes at RT. Samples were placed on the magnetic rack, and the supernatant containing purified DNA was collected. Presence of the DC was quantified by RT-qPCR of the target genes *ribokinase* (*FOXG_13522*)^3^ or the *pg1* gene (*FOXG_14695*) located on this chromosome, using primer combinations ribokinase qFor/ribokinase qRev or pg1 ORF F/pg1 ORF R, respectively. The *actin* gene (*FOXG_01569*) located on core chromosome 5 was used as a reference (primers Act q7/Act q8).

Three technical replicates were performed for each biological sample. The relative proportion of nuclei carrying the DC in the population was quantified using the Pfaffl method^50^, where DNA levels of the DC-specific genes were normalized to those of the *actin* reference gene, and normalized against that in the initial population at time zero (t0), considering primer efficiencies. A relative ratio of 1 represents 100% retention of the DC. The following formula was used:

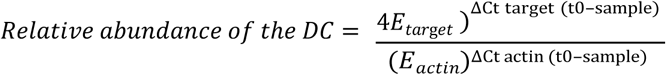

where target represents either the *ribokinase* gene or *pg1* gene, and E represent the amplification efficiency for each amplicon. Primer efficiencies were empirically determined from standard curves, where E is defined as (%*Efficiency*/100) + 1 (Supplementary Table S4).

Flow cytometry data collection from samples was performed on a BD FACSCelesta flow cytometer with BD FACSDiva Software version 9.0 (Becton Dickinson Biosciences, Franklin Lakes, NJ, United States) as described^17^. To ensure the accuracy of the measurements, two successive gatings steps were applied to consider only viable single-cell events for data acquisition. First, cell viability staining was performed by adding 20 µM propidium iodide directly to the sample followed by 5 minutes incubation in the dark. Second, cell multiplets were identified and excluded from the final dataset. Following data acquisition, the FlowJo v10.10 software (Tree Star, OR, USA) was used to analyse cell populations. Viable single-cell populations were down-sampled to 100’000 events per sample. Presence of the DC in the population was inferred by detection of the fluorescent signal from *3xclover* (491 nm excitation, 516 nm detection). The *3xclover* parental strain from a given passaging experiment and the unlabelled wild type (wt) strain served as positive and negative controls, respectively.

### Competition experiments

For *in vitro* competition experiments, PDA plates or MMA plates containing sucrose or ribose as the sole carbon source were spot-inoculated with a 1:1 mix containing 5×10^5^ freshly obtained microconidia from each strain. Plates inoculated with 10^6^ microconidia from each strain separately were used as non-competition controls. After 3 days (for flow cytometry analysis) or 4 days (for RT-qPCR analysis) incubation, total microconidia were collected from the colony and stored at -80°C. Three and five biological replicates were performed for flow cytometry and RT-qPCR analysis, respectively.

For competition assays using tomato plants as nutrient source, seedlings of *Solanum lycopersicum* cv. Moneymaker were maintained in a growth chamber as described^51^. After 28 days, plants were harvested, killed by submerging in liquid nitrogen for 1 minute, placed in a sterile petri dish, and inoculated by equally distributing ten 5 μl drops containing a 1:1 mix (2.5×10^6^ microconidia/ml each) of strains MRL8996 and MRL8996 -DC10, or 5×10^6^ microconidia/ml of MRL8996 alone as a control along every individual plant. After 7 days of incubation at 28°C and 100% humidity, the biological material was collected, freeze-dried and lyophilized for DNA extraction and RT-qPCR analysis as described above. Five biological replicates were performed.

### Determination of relative fitness of fluorescent versus non-fluorescent strains by flow cytometry

Frozen microconidia stocks from the initial 1:1 inoculation mixture and from the post-competition population were re-grown 3 days in PDB at 28 °C and 170 rpm. Microconidia were collected and quantified as described above. Aliquots of 5×10^6^ spores were resuspended in 500 μl of 50 mM sodium citrate (pH= 7.1) and analysed by flow cytometry as described above. To prevent sample size bias and ensure equal statistical weighting, viable single-cell populations were uniformly down-sampled before data extraction. The down-sampling threshold was maximized independently for each experimental block, based on its lowest common viable event count (60,000 events for Fol4287, 100,000 events for MRL8996 competition experiments).

Relative fitness (W) was calculated using the formula: 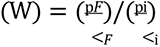 Where *p_i_* and *q_i_* represent the proportion of fluorescent and non-fluorescent microconidia, respectively, in the initial inoculum (usually around 0.5), while *p_f_* and *q_f_* represent the proportions of fluorescent and non-fluorescent microconidia, respectively, in the post-competition population.

### Determination of relative fitness of MRL8996 wild type versus MRL8996 -DC10 strain by qPCR

The proportion of the MRL8996 wild type strain carrying DC10 (*p*) in conidia of the initial inoculum mixture (*pi*) or from the post-competition population (*pf*) was determined by qPCR of total genomic DNA. Calculations were performed using the Δ*Ct* values of the *pg1* gene and the reference gene (*actin*), normalized to a time 0 control sample containing 100% wild-type MRL8996 cells. The absolute proportion of the wild type strain (*p*) was calculated using the Pfaffl method ^50^ to adjust for primer efficiencies (Supplementary Table S4). The proportion of strain MRL8996 -DC10 (*q*) was calculated as follows: *q* = 1 − *p*.

### Data correction models for DC loss

The corrected relative fitness coefficient (*W*_corr_) was calculated to decouple the metabolic fitness cost in the *in vitro* competition experiments from the basal DC loss rate (λ), measured in a non-competitive pure culture. The corrected proportions of cells in the post-competition population (*p_f_*) carrying (*p*>*f*) or lacking the DC (*q*>*f*) is defined as:

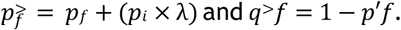

The corrected relative fitness (*W*_corr_) is expressed as 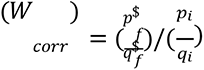 The λ correction factor was not required for *in-planta* assays, since non-competition controls demonstrated stable maintenance of DC10 (λ ≈ 0). Furthermore, competitive exclusion *in planta* occasionally resulted in the outcompeted strain falling below the qPCR detection limit. In such cases approaching complete population fixation, technical variance could result in calculated *pf* ≥ 1 (with q*f* = 0). To avoid a zero denominator in the fitness equation, *pf* was adjusted to a maximum detection limit of *pf* = 0.99 (with q*f* = 0.01) to accurately reflect the dominance of the DC10-carrying strain.

### Data and code availability

The raw genome sequencing reads of the evolved Fol4287 and MRL8996 lines are deposited in NCBI SRA under project PRJNA1522590 (Accession: SRR40492194-202). The genome sequencing reads for the MRL8996 -DC10 strain are deposited in NCBI SRA (Accession: SRR40443837). Scripts and R codes used for the extended figures and all the source files are available at https://github.com/doddipetto/spok-source-data-and-figures.

## Acknowledgements

We are grateful to Prof. Aaron Vogan, Uppsala University, Sweden, for stimulating discussions on *spoks* and *Starships* and for sharing unpublished information; and to Prof. Daniel Croll, University of Neuchâtel, Switzerland, for hosting G.P.P. during a 3-month predoctoral stay. We thank Mariló Alcaide Caballero, Universidad de Córdoba, for excellent technical assistance.

## Author contributions

Conceptualization: G.P.P., A.D., D.H.A., P.G.E., L.J.M., M.S.L.B. and ADP. Formal analysis: G.P.P., A.D., D.H.A., P.G.E., L.J.M., M.S.L.B. and ADP. Funding acquisition: P.G.E., L.J.M., M.S.L.B. and ADP. Investigation: G.P.P., A.D., D.H.A., P.G.E., L.J.M., M.S.L.B. and ADP. Methodology: G.P.P., A.D., D.H.A., P.G.E., L.J.M., M.S.L.B. and ADP. Project administration: P.G.E., L.J.M., M.S.L.B. and ADP. Resources: L.J.M., M.S.L.B. and A.D.P. Software: G.P.P., A.D., D.H.A., P.G.E., L.J.M., M.S.L.B. and ADP. Supervision: P.G.E., L.J.M., M.S.L.B. and ADP. Validation: G.P.P., A.D., D.H.A., P.G.E., L.J.M., M.S.L.B. and ADP. Visualization: G.P.P., A.D., D.H.A., M.S.L.B. and ADP. Writing (original draft): G.P.P. and A.D.P. Writing (review and editing): G.P.P., A.D., D.H.A., P.G.E., L.J.M., M.S.L.B. and ADP.

## Supplementary information

**Supplementary Table 1.** Fungal strains used in the study.

**Supplementary Table 2.** Plasmids used in the study.

**Supplementary Table 3.** Oligonucleotides used.

**Supplementary Table 4**. Efficiency of qPCR.

**Supplementary Table 5.** Pairwise alignment coverage and mean nucleotide identity between the chromosomes of *F. oxysporum* MRL8996 and *F. oxysporum* f. sp. *lycopersici* 4287.

**Supplementary Table 6.** Chromosome correspondence between Fol4287 and MRL8996.

**Supplementary Table 7**. Tip metadata of the chromosome-10 control tree (464 tips).

**Supplementary Table 8**. Whole-genome alignment statistics of all screened Fusarium assemblies against Fol4287 chromosomes 12 and 10.

**Supplementary Table 9.** tblastn hits of the Fol4287 Spok protein (XP_018254443.1) against the extracted chromosome-12 intervals of all genomes.

**Supplementary Table 10**. Individual blastn alignment fragments of the Fol4287 *Starship* reference against each genome, before merging into per-genome coverage.

**Supplementary Table 11**. *Starship*, *captain* and *spok* status per genome.

**Supplementary Table 12.** Detailed *Starship, captain* and *spok* detection metrics per genome.

**Supplementary Table 13**. Per-genome *spok* presence calls on the chromosome-12 homolog.

**Supplementary Table 14**. Gene ontology categories significantly enriched on the dispensable core chromosome of MRL8996 (DC10).

**Supplementary Table 15.** Genome-wide gene ontology enrichment per chromosome of MRL8996.

**Supplementary Table 16**. Normalised read coverage of MRL8996 wild type and MRL8996 -DC10 in 10 kb windows.

**Supplementary Table 17**. Per-chromosome coverage and copy number in MRL8996 wild type and MRL8996 -DC10.

---

**Extended Data Figure 1.**
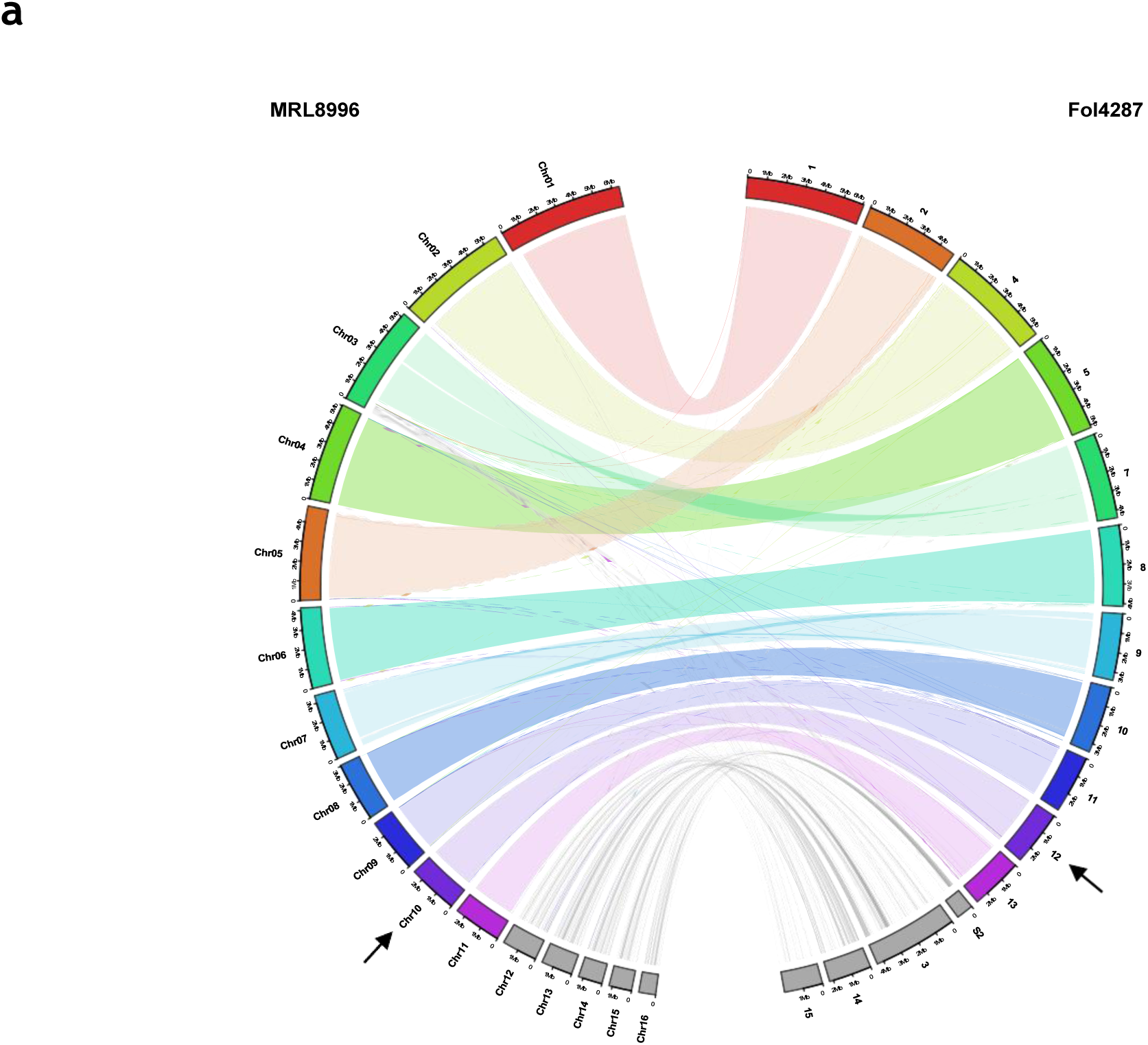
Synteny of chromosomes between *F. oxysporum* isolates MRL8996 and Fol4287. **a.** Circos plot of whole-genome minimap2 alignments between the genome assemblies of isolates Fol4287 (GCA_030020275.1) and MRL8996 (GCA_009746015.2). Lines connecting chromosomes of the two genomes represent homologous regions (>5 kb). Core chromosomes are in color, accessory chromosomes/regions are in grey. Note that the dispensable core chromosome 12 in Fol4287 is syntenic to MRL8996 chromosome 10 (arrows).

**Extended Data Figure 2.**
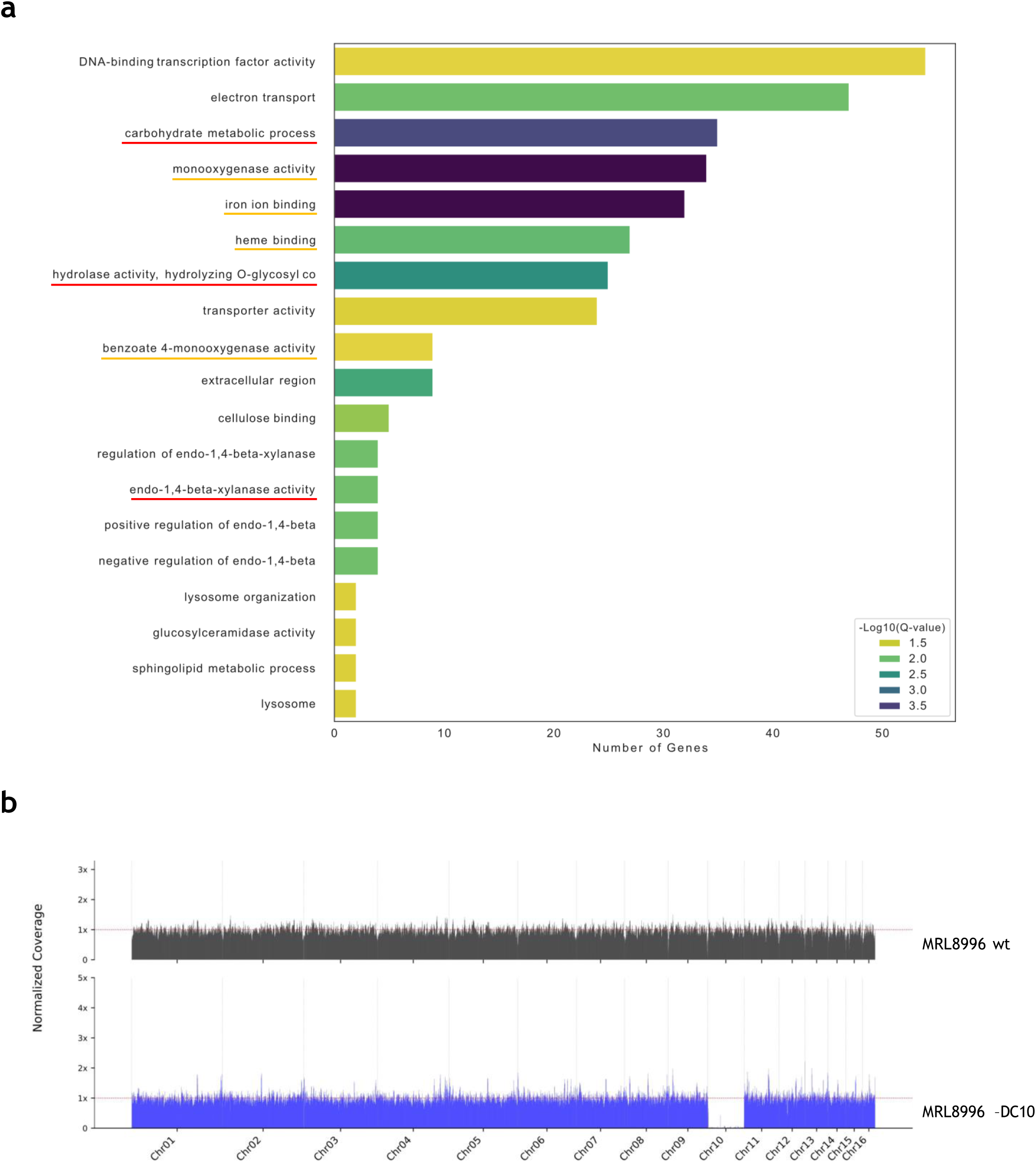
Dispensable core chromosome 10 is enriched in genes related to metabolism of non-preferred carbon sources and defensive stress responses. **a.** Gene ontology analysis of functional categories enriched on DC10 from MRL8996. Enrichment of each GO term was tested with a one-sided Fisher exacttest comparing the annotated genes on DC10 with all remaining annotated genes of the MRL8996 genome; only categories significant after Benjamini-Hochberg correction (q < 0.05) are shown. Colours indicate the FDR-corrected significance (-log10 q). Categories related with the use of non-preferred carbon sources or with defensive stress responses/xenobiotic degradation are underlined in red and orange, respectively. **b.** Normalised read coverage distribution (10 kb windows) from Nanopore sequencing of the MRL8996 wild type strain and of a monosporic isolate that has spontaneously lost DC10 (MRL8996 –DC10). Relative positions of the different chromosomes are indicated.

**Extended Data Figure 3.**
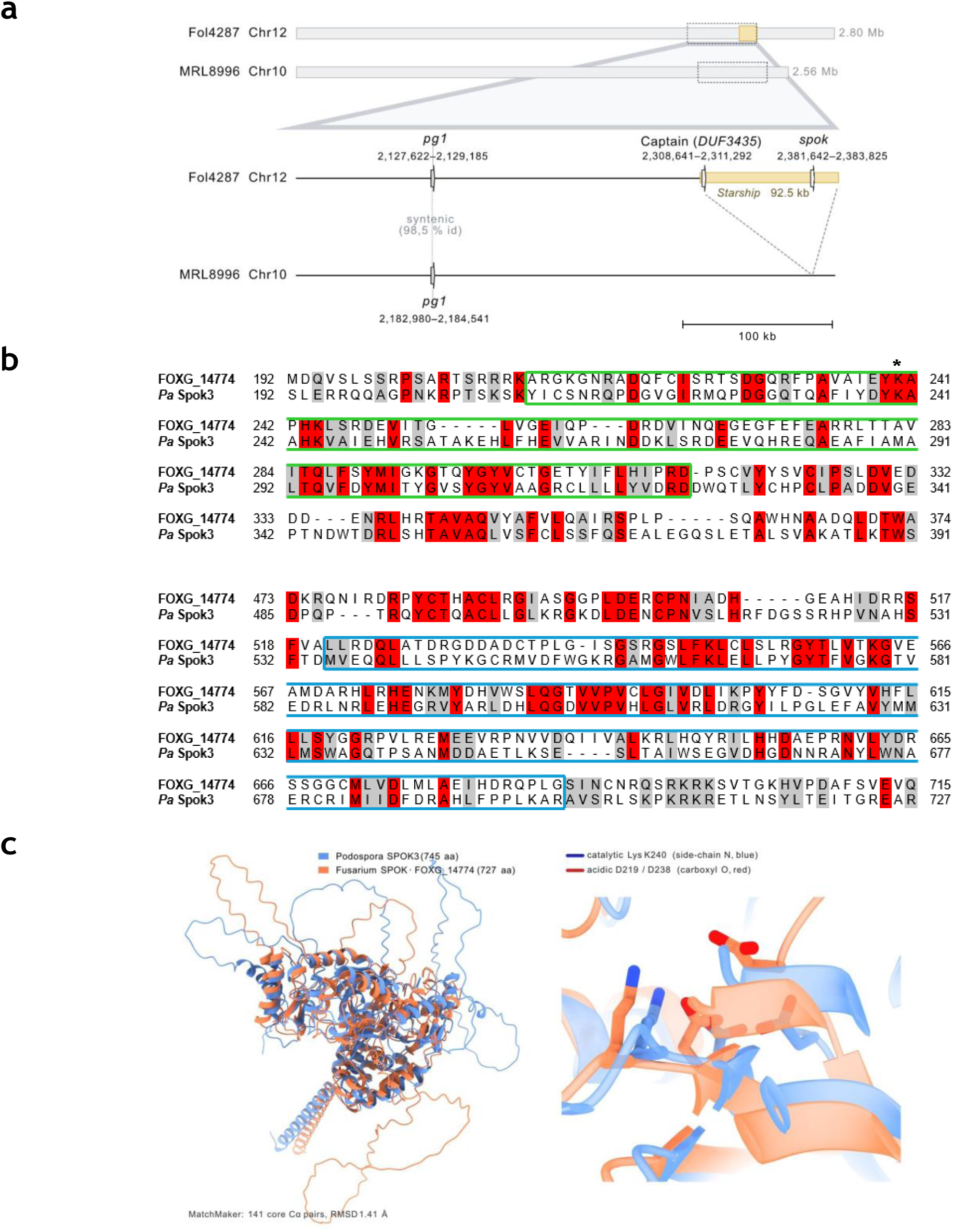
Chromosome 12 of Fol4287 carries a *Starship* element harbouring a *spok* gene. **a.** Fol4287 chromosome 12 carries an insertion of a *Starship* giant transposon (yellow box) harbouring a spore killer (*spok*) gene, both of which are absent the homologous chromosome 10 of MRL8996. Upper diagrams show entire chromosomes, lower diagrams show an amplification of the indicated chromosomal region. Positions of the *DUF3435 captain* and the *spok* genes in *Starship* are indicated. **b.** Alignment of the predicted product of the Fol4287 *spok* gene (FOXG_14774) with the nuclease (green box) and the kinase (blue box) domains of Spok3 from *Podospora anserina*. Conserved and similar residues are shaded in red and grey, respectively. The conserved lysin residue at position 240, which is essential for Spok killing activity, is marked with an asterisk. **c.** AlphaFold2 protein structure predictions of *F. oxysporum spok FOXG_14774* (XP_018254443.1) and *P. anserina* SPOK3 (UniProt A0A516F180) were aligned using ChimeraX16 MatchMaker. Right panel shows the positions of the K240 catalytic and the D219/D238 DNA-binding residues of the killing domain.

**Extended Data Figure 4.**
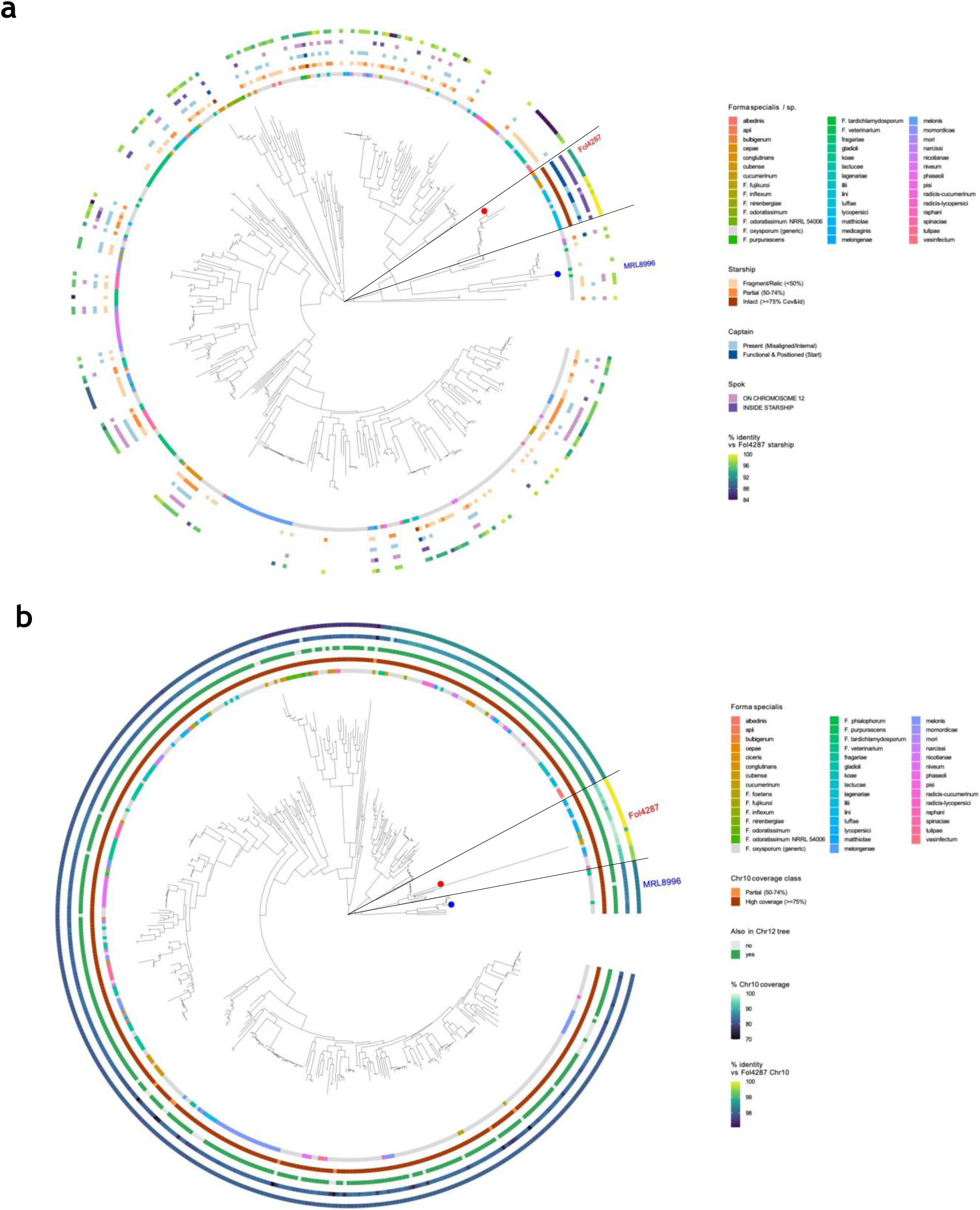
Phylograms of Fol4287 core chromosomes 12 and 10 in *Fusarium*. Maximum-likelihood trees (midpoint-rooted) of DC12 (RefSeq NC_030997.1) (**a**) and of chromosome 10 (RefSeq NC_030995.1) (**b**) from Fol4287. Among the 1,252 publicly available *Fusarium* genome assemblies, obtained with ultrafast bootstrap (n = 1,000) approximations from IQ-tree. Assemblies recovering less than 70% of the reference chromosome were excluded from all downstream phylogenetic analyses; 421 assemblies passed this filter for chromosome 12, while 468 passed for chromosome 10. **a.** Color scales of circles (from inner to outer) denote *Fusarium* species or *F. oxysporum* forma specialis; presence/absence of *Starship*, *DUF3435 captain,* and *spok*, respectively; and % identity with the Fol4287 *Starship*. Note that intact *Starships* including a *spok* gene are present on chromosome 12 homologs of closely related *F. oxysporum* isolates (delimited by lines). **b.** Circles (from inner to outer) indicate *Fusarium* species or *F. oxysporum* forma specialis; chromosome 10 coverage class; presence/absence in the reciprocal tree of Fol4287 chromosome 12 homologs; % coverage of chromosome 10; and mean nucleotide identity with Fol4287 chromosome 10. Positions of Fol4287 and MRL8996 on trees are indicated.

**Extended Data Figure 5.**
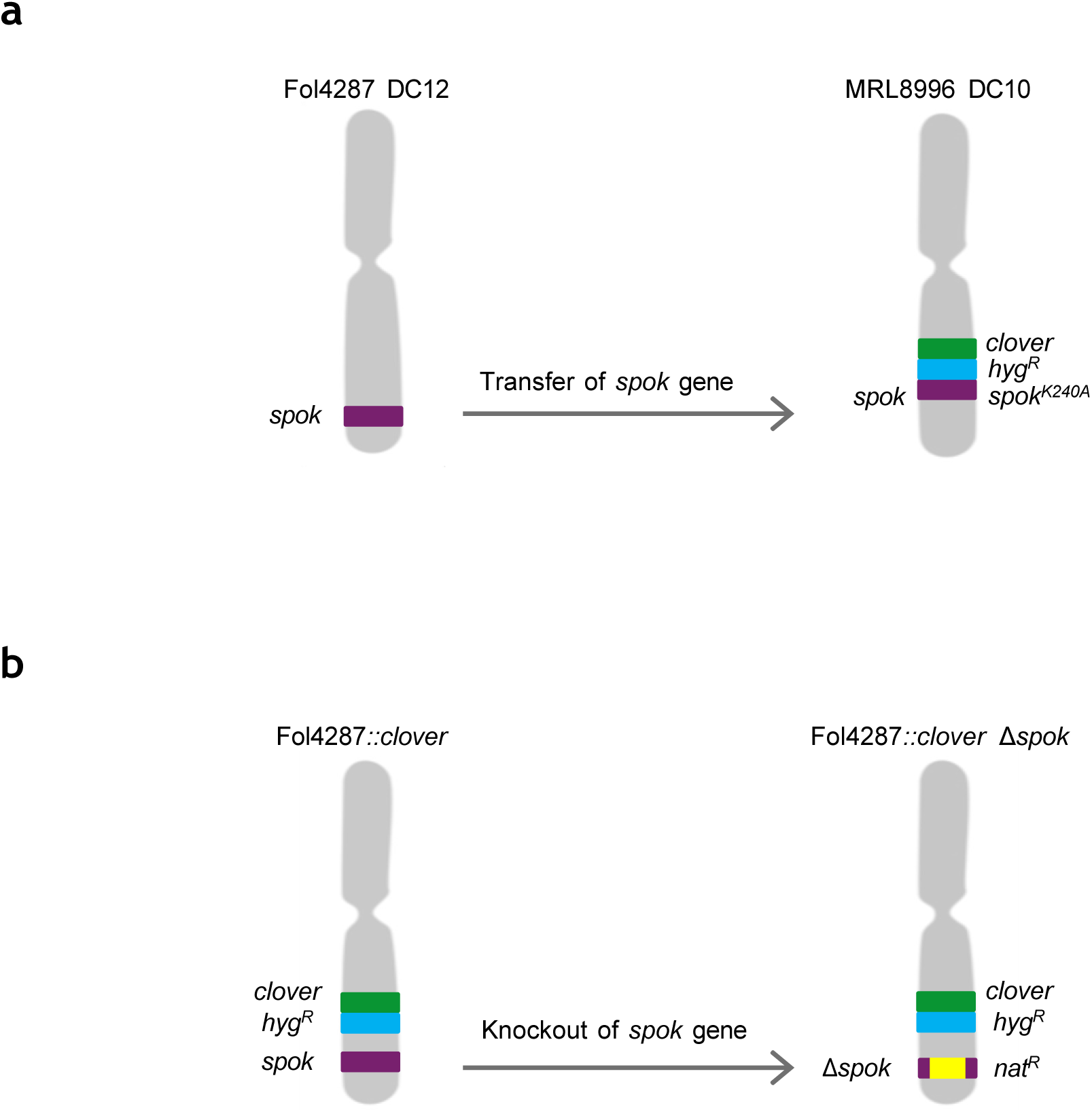
Two complementary approaches to test the role of *spok* in maintenance of DC10/DC12. **a.** Transfer of the *spok* gene or a killing-deficient *spok^K240A^* mutant allele thereof, from Fol4287 DC12 to MRL8996 DC10. **b.** Targeted replacement of the *spok* gene on Fol4287 chromosome 12 with the nourseothricin resistance gene (*nat^R^*). In both approaches, the green fluorescent protein gene *3xclover* and the hygromycin resistance cassette (*hyg^R^*), were inserted in the DC to allow quantification of spontaneous loss (see Methods section).

**Extended Data Figure 6.**
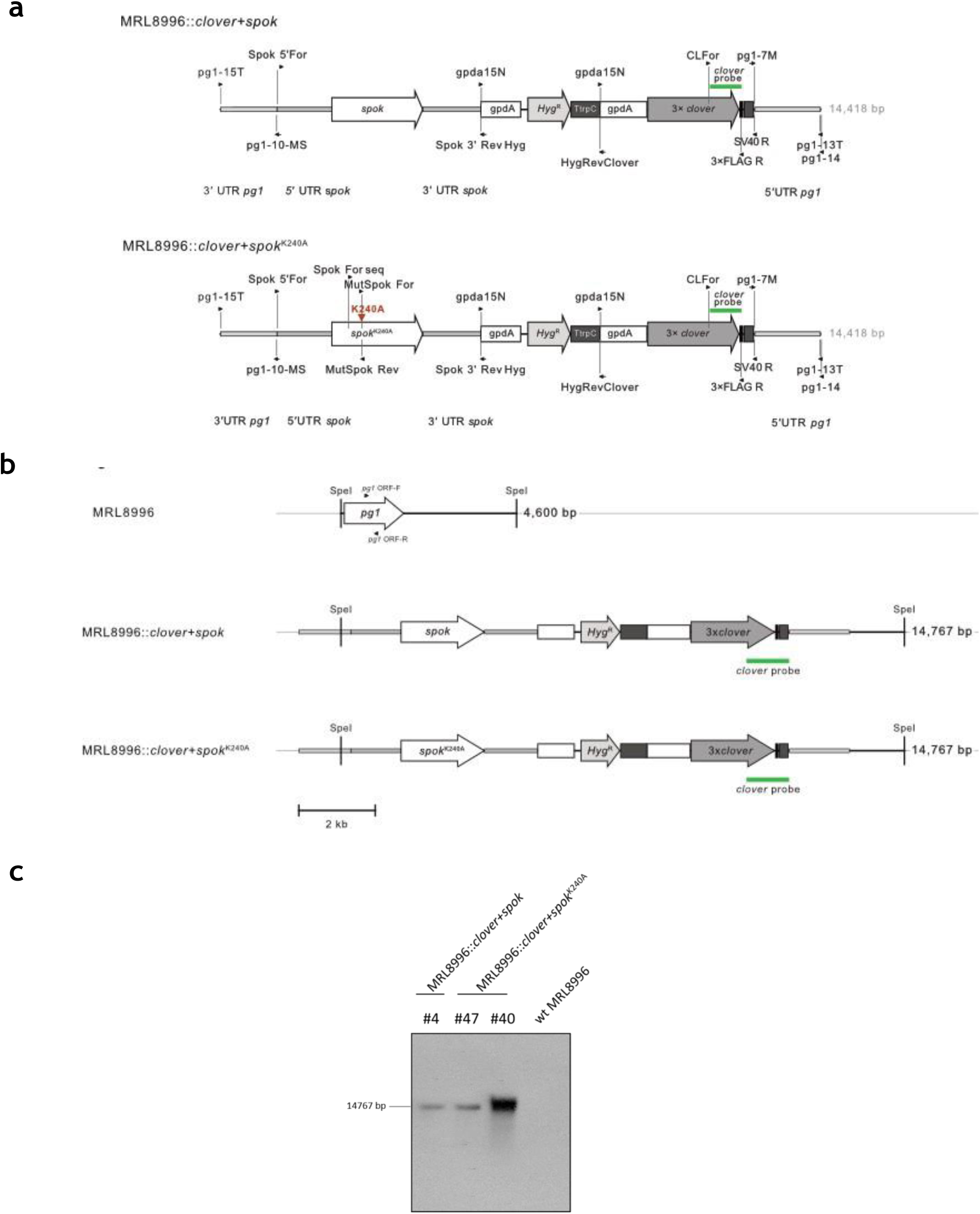
Generation of MRL8996::*clover+spok* and MRL8996::*clover+spok*^K240A^ strains. **a.** Schematic representation of the MRL8996::*clover+spok* and MRL8996::*clover+spok*^K240A^ constructs with relative positions of the flanks for targeted replacement of the *pg1* locus on DC10, the probe for Southern blot, and relevant primers. **b.** Physical map of the obtained strains with relative positions of restriction sites and probes used for Southern blot. Predicted band sizes are on the right. **c.** Southern blot analysis of the obtained transformants. Genomic DNA of the wild type (wt) and the PCR screening-positive transformants was treated with *SpeI,* separated on a 0,7% agarose gel, transferred to a nylon membrane, and hybridized with the *clover* probe (see **a**). Molecular size of the hybridizing band is indicated on the left. Transformants #4 of MRL8996::*clover+spok* and #47 of MRL8996::*clover+spok*^K240A^ show hybridizing bands consistent with homologous replacement of the *pg1* locus on DC10 with the construct from (**a**).

**Extended Data Figure 7.**
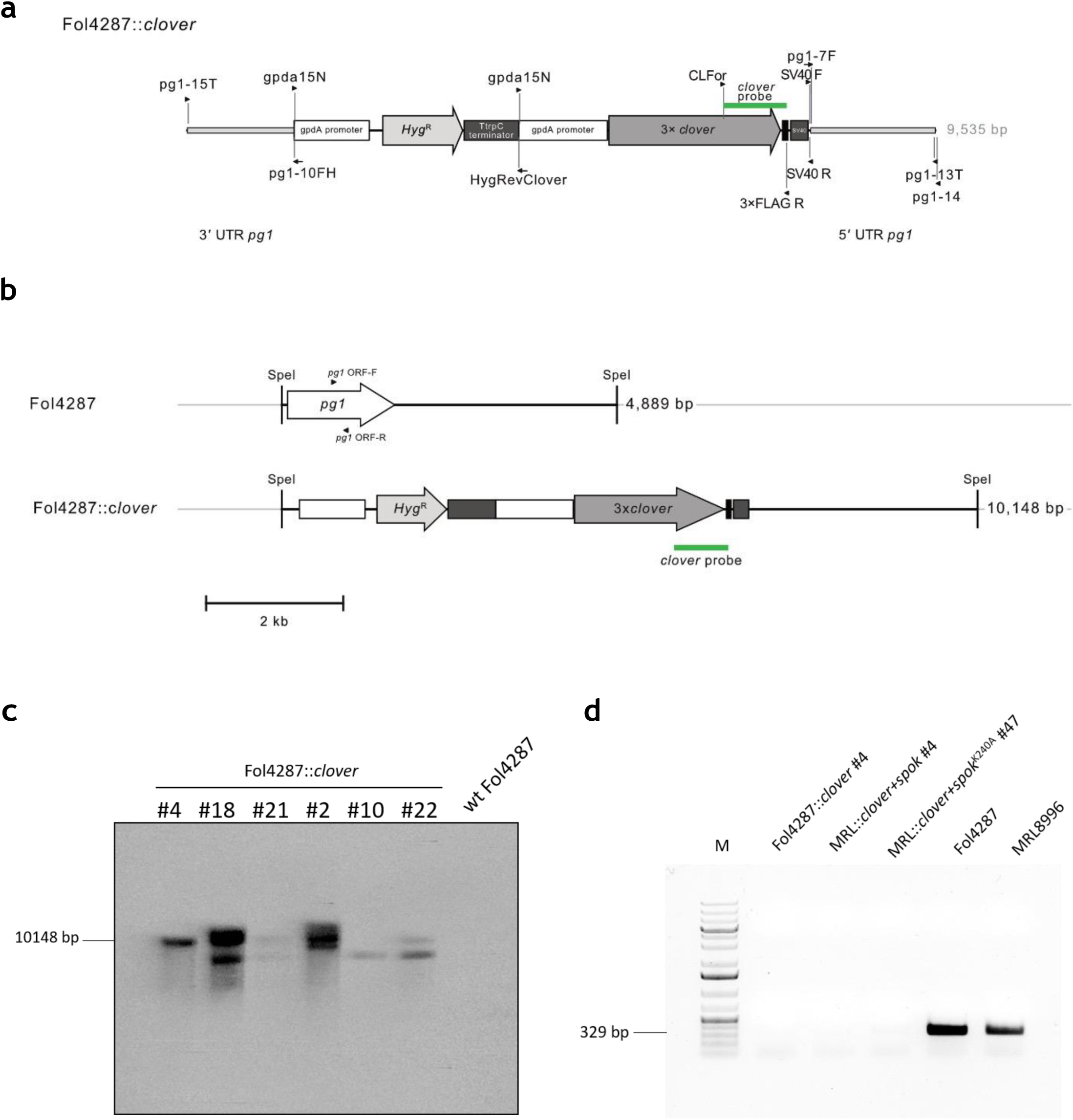
Generation of the Fol4287::*clover* strain. **a.** Schematic representation of the Fol4287::*clover* construct with relative positions of the flanks for targeted replacement of the *pg1* locus on DC12, the probe for Southern blot, and relevant primers. **b.** Physical map of the *pg1* locus in the Fol4287 wild type and Fol4287::*clover* insertion mutant. with relative positions of restriction sites and probes used for Southern blot, and relevant primers. Predicted band sizes are on the right. **c.** Southern blot analysis of obtained transformants. Genomic DNA of the wild type (wt) and independent PCR screening-positive transformants was treated with *SpeI,* separated on a 0,7% agarose gel, transferred to a nylon membrane, and hybridized with the *clover* probe (see **a**). Molecular size of the hybridizing band is indicated on the left. Transformant #4 shows a hybridizing band consistent with homologous replacement of the *pg1* locus on DC12 with the construct from (**a**). **d.** Agarose gel electrophoresis of PCR products obtained using the primer pair *pg1* ORF-F and *pg1* ORF-R (indicated in **b**) with genomic DNA extracted from the indicated strains. M, molecular size marker. Absence in the selected mutants, of a 329 bp amplification band detected in the two wild type strains indicates homologous insertion at the *pg1* locus.

**Extended Data Figure 8.**
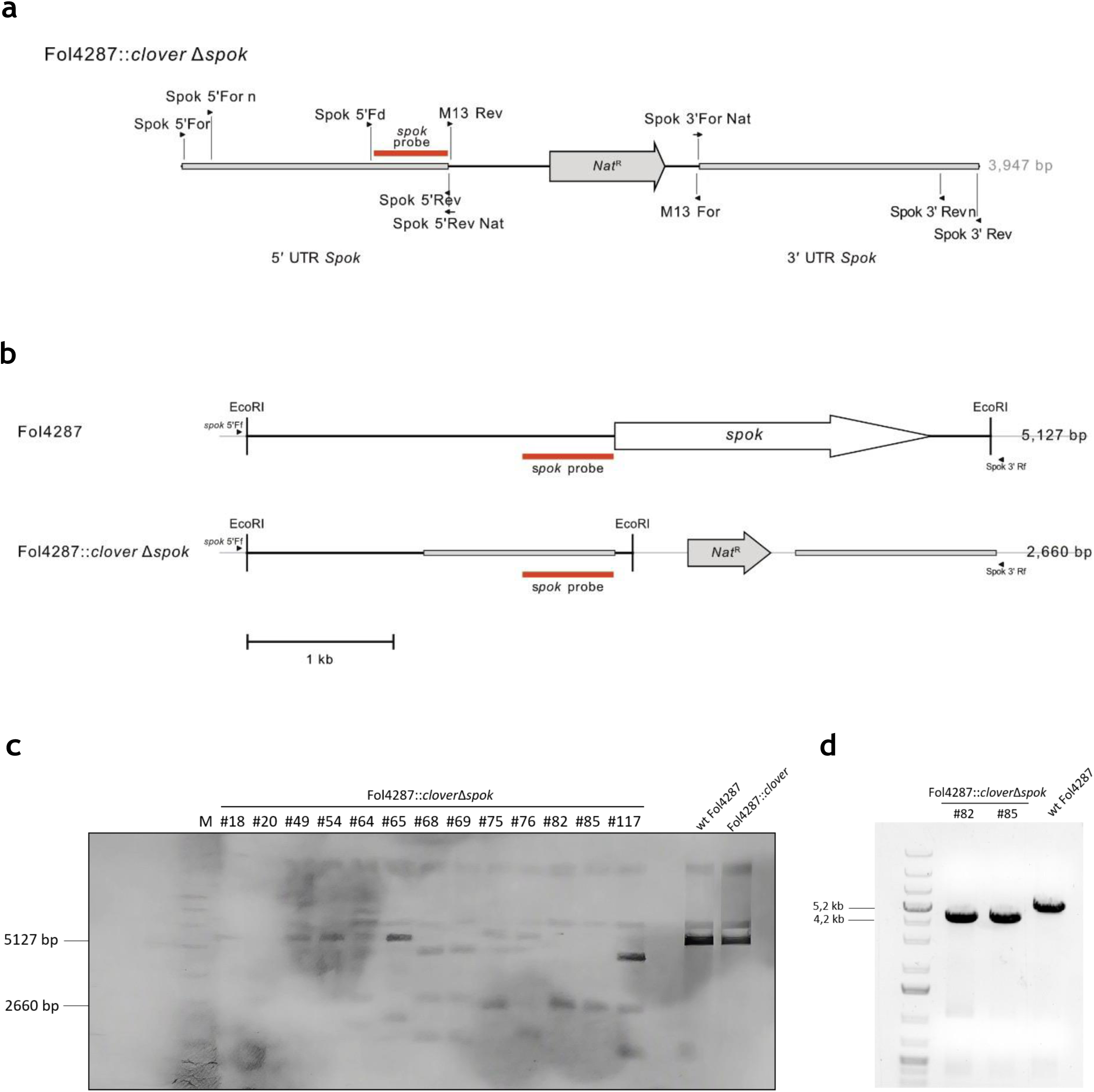
Generation of Fol4287::*clover*Δ*spok* knockout mutants. **a.** Schematic representation of the Fol4287Δ*spok* construct with relative positions of the flanks for targeted replacement of the of the *spok* gene with the *Nat^R^* resistance cassette, the probe for Southern blot, and relevant primers. **b.** Physical map of the *spok* locus in the Fol4287 wild type and Fol4287Δ*spok* knockout mutant. with relative positions of restriction sites and probes used for Southern blot, and relevant primers. Predicted band sizes are on the right. **c.** Southern blot analysis of obtained transformants. Genomic DNA of the wild type (wt) and independent PCR screening-positive transformants was treated with *EcoRI,* separated on a 0,7% agarose gel, transferred to a nylon membrane, and hybridized with the *spok* probe (see **a**). Molecular size of the hybridizing band is indicated on the left. Transformants #82 and #85 (named hereafter Fol4287::*clover* Δ*spok* #1 and #2, respectively) show hybridizing bands consistent with homologous replacement of the *spok* gene with the construct from (**a**). **d.** agarose gel electrophoresis of PCR products obtained using the primer pair *spok* 5 Ff and *spok* 3 Rf (indicated in **b**) with genomic DNA extracted from the indicated strains. M, molecular size marker. Presence in the Δ*spok* mutants, of a 4.2 kb amplification band instead of the 5.2 kb band of the wild type strain indicates homologous replacement of the *spok* locus with the *Nat^R^* resistance cassette.

**Extended Data Figure 9.**
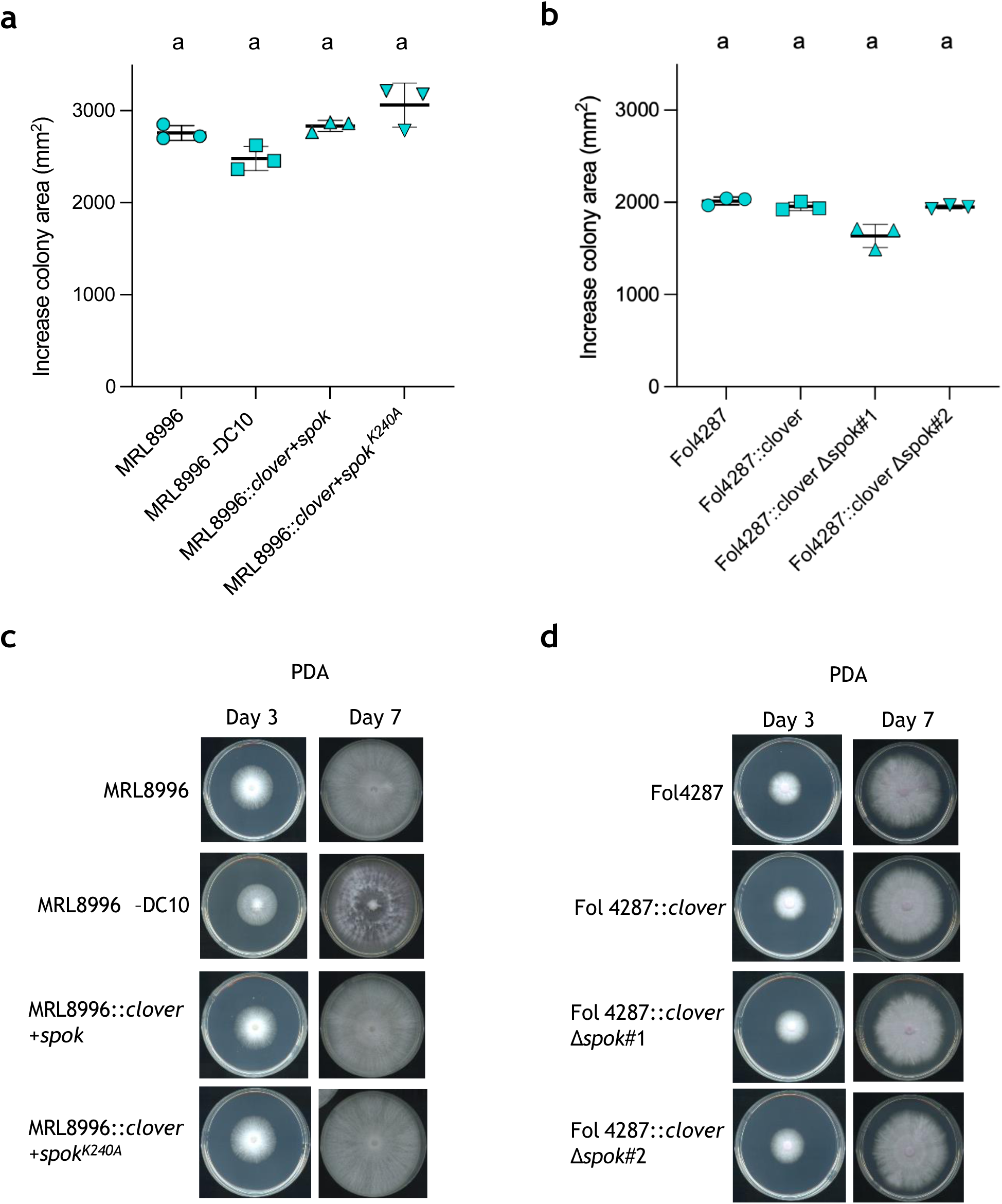
Presence/absence of *spok* does not affect colony growth rate. **a, b.** Growth rates of the MRL8996 (**a**) and Fol4287 (**b**) wild type and derived strains harbouring the indicated *spok* alleles on DC10/12. A total of 10^6^ microconidia were spot-inoculated at the center of a PDA plate and incubated at 28°C. Plates were scanned at different time points. Growth rate was calculated as increase in colony area between day 3 and 7. Each data point represents an independent plate. Error bars, SD. In each graph, strains with the same letter do not differ significantly according to two-way ANOVA followed by Dunnett’s multiple comparison test (p>0.05). **c, d.** Representative images of colonies of the MRL8996 (**c**) and Fol4287 (**d**) wild type and derived strains grown on PDA for 3 days.

## References

1. Ferree, P. M., et al. What is a B chromosome? Early definitions revisited. G3 Genes|Genomes|Genetics 14, doi:10.1093/g3journal/jkae068 (2024).

2 Komluski, J., Stukenbrock, E. H. & Habig, M. Non-Mendelian transmission of accessory chromosomes in fungi. Chromosome Res 30, 241–253, doi:10.1007/s10577-022-09691-8 (2022).

3 Vlaardingerbroek, I., Beerens, B., Schmidt, S. M., Cornelissen, B. J. & Rep, M. Dispensable chromosomes in Fusarium oxysporum f. sp. lycopersici. Mol Plant Pathol 17, 1455–1466, doi:10.1111/mpp.12440 (2016).

4 Dijkstra, J. et al. Accessory Chromosome Contributes to Virulence of Banana Infecting Fusarium oxysporum Tropical Race 4. Molecular Plant Pathology 26, e70146, 10.1111/mpp.70146 (2025).

5 Croll, D. & McDonald, B. A. The accessory genome as a cradle for adaptive evolution in pathogens. PLoS Pathog 8, e1002608, doi:10.1371/journal.ppat.1002608 (2012).

6 Yang, H., Yu, H. & Ma, L. J. Accessory Chromosomes in Fusarium oxysporum. Phytopathology 110, 1488–1496, doi:10.1094/PHYTO-03-20-0069-IA (2020).

7 Habig, M., Quade, J. & Stukenbrock Eva, H. Forward Genetics Approach Reveals Host Genotype-Dependent Importance of Accessory Chromosomes in the Fungal Wheat Pathogen Zymoseptoria tritici. mBio 8, 10.1128/mbio.01919-01917, doi:10.1128/mbio.01919-17 (2017).

8 MacLean, R. C. & San Millan, A. Microbial Evolution: Towards Resolving the Plasmid Paradox. Curr Biol 25, R764–767, doi:10.1016/j.cub.2015.07.006 (2015).

9 Dean, R. et al. The Top 10 fungal pathogens in molecular plant pathology. Mol Plant Pathol 13, 414–430, doi:10.1111/j.1364-3703.2011.00783.x (2012).

10 Ortoneda, M. et al. Fusarium oxysporum as a Multihost Model for the Genetic Dissection of Fungal Virulence in Plants and Mammals. Infection and Immunity 72, 1760–1766, doi:10.1128/iai.72.3.1760-1766.2004 (2004).

11 Nucci, M. & Anaissie, E. Invasive fusariosis. Clinical Microbiology Reviews 36, e00159–00122, doi:10.1128/cmr.00159-22 (2023).

12 Zhang, Y. et al. The genome of opportunistic fungal pathogen Fusarium oxysporum carries a unique set of lineage-specific chromosomes. Commun Biol 3, 50, doi:10.1038/s42003-020-0770-2 (2020).

13 Bravo Nunez, M. A., Lange, J. J. & Zanders, S. E. A suppressor of a wtf poison-antidote meiotic driver acts via mimicry of the driver’s antidote. PLoS Genet 14, e1007836, doi:10.1371/journal.pgen.1007836 (2018).

14 Nuckolls, N. L. et al. S. pombe wtf drivers use dual transcriptional regulation and selective protein exclusion from spores to cause meiotic drive. PLOS Genetics 18, e1009847, doi:10.1371/journal.pgen.1009847 (2022).

15 Vogan, A. A. et al. Combinations of Spok genes create multiple meiotic drivers in Podospora. Elife 8, doi:10.7554/eLife.46454 (2019).

16 Sandell, L. et al. The role of toxin/antidote genes in the maintenance and evolution of accessory chromosomes in Fusarium. Genetics 231, doi:10.1093/genetics/iyaf197 (2025).

17 López Díaz, C., et al. Transposons and accessory genes drive adaptation in a clonally evolving fungal pathogen. Nature Communications 16, 6982, doi:10.1038/s41467-025-62213-y (2025).

18 Ma, L. J. et al. Comparative genomics reveals mobile pathogenicity chromosomes in Fusarium. Nature 464, 367–373, doi:10.1038/nature08850 (2010).

19 Chang, D. C. et al. Multistate Outbreak of Fusarium Keratitis Associated With Use of a Contact Lens Solution. JAMA 296, 953–963, doi:10.1001/jama.296.8.953 (2006).

20 Doddi, A., Puebla Planas, G., López-Berges, M. S. & Di Pietro, A. A near-complete genome assembly of the Fusarium oxysporum keratitis isolate MRL8996. bioRxiv, 2026.2007.2006.736765, doi:10.64898/2026.07.06.736765 (2026).

21 Gluck-Thaler, E. et al. Giant Starship Elements Mobilize Accessory Genes in Fungal Genomes. Mol Biol Evol 39, doi:10.1093/molbev/msac109 (2022).

22 Zanders, S. & Johannesson, H. Molecular Mechanisms and Evolutionary Consequences of Spore Killers in Ascomycetes. Microbiol Mol Biol Rev 85, e0001621, doi:10.1128/MMBR.00016-21 (2021).

23 Fischer, M. S. & Glass, N. L. Communicate and Fuse: How Filamentous Fungi Establish and Maintain an Interconnected Mycelial Network. Frontiers in Microbiology Volume 10 - 2019, doi:10.3389/fmicb.2019.00619 (2019).

24 Rosales, R. C. P. & Pietro, A. D. Vegetative Hyphal Fusion Is Not Essential for Plant Infection by Fusarium oxysporum. Eukaryotic Cell 7, 162–171, doi:10.1128/ec.00258-07 (2008).

25 Palos-Fernández, R., Turrà, D. & Pietro, A. D. The Gal4-Type Transcription Factor Pro1 Integrates Inputs from Two Different MAPK Cascades to Regulate Development in the Fungal Pathogen Fusarium oxysporum. Journal of Fungi 8, 1242 (2022).

26 Gerdes, K., Rasmussen, P. B. & Molin, S. Unique type of plasmid maintenance function: postsegregational killing of plasmid-free cells. Proceedings of the National Academy of Sciences 83, 3116–3120, doi:10.1073/pnas.83.10.3116 (1986).

27 Yang, Q. E. & Walsh, T. R. Toxin–antitoxin systems and their role in disseminating and maintaining antimicrobial resistance. FEMS Microbiology Reviews 41, 343–353, doi:10.1093/femsre/fux006 (2017).

28 Ayhan, D. H. et al. Host-specific adaptation in Fusarium oxysporum correlates with distinct accessory chromosome content in human and plant pathogenic strains. mBio 16, e00951–00925, doi:10.1128/mbio.00951-25 (2025).

29 Vogan, A. A. et al. The Enterprise, a massive transposon carrying Spok meiotic drive genes. Genome Res 31, 789–798, doi:10.1101/gr.267609.120 (2021).

30 Lai, E. C. & Vogan, A. A. Proliferation and dissemination of killer meiotic drive loci. Curr Opin Genet Dev 82, 102100, doi:10.1016/j.gde.2023.102100 (2023).

31 Like Fokkens, S. S., Lanelle R. Connolly, Remco Stam, Sarah M. Schmidt, Kristina M. Smith, Michael Freitag, Martijn Rep. The multi-speed genome of Fusarium oxysporum reveals association of histone modifications with sequence divergence and footprints of past horizontal chromosome transfer events. 10.1101/465070 (2018).

32 Bastiaans, E., Debets, A. J. M. & Aanen, D. K. Experimental evolution reveals that high relatedness protects multicellular cooperation from cheaters. Nature Communications 7, 11435, doi:10.1038/ncomms11435 (2016).

33 Debets, A. J. M. & Griffiths, A. J. F. Polymorphism of het-genes prevents resource plundering in Neurospora crassa. Mycological Research 102, 1343–1349, 10.1017/S095375629800639X (1998).

34 Glass, N. L. & Kaneko, I. Fatal Attraction: Nonself Recognition and Heterokaryon Incompatibility in Filamentous Fungi. Eukaryotic Cell 2, 1–8, doi:10.1128/ec.2.1.1-8.2003 (2003).

35 Vangalis, V., Likhotkin, I., Knop, M., Typas, M. A. & Papaioannou, I. A. Starvation-induced cell fusion and heterokaryosis frequently escape imperfect allorecognition systems in an asexual fungal pathogen. BMC Biology 19, 169, doi:10.1186/s12915-021-01101-5 (2021).

36 Ishikawa, F. H. et al. Heterokaryon Incompatibility Is Suppressed Following Conidial Anastomosis Tube Fusion in a Fungal Plant Pathogen. PLOS ONE 7, e31175, doi:10.1371/journal.pone.0031175 (2012).

37 Ament-Velásquez, S. L. & Saupe, S. J. NOD-Like Receptor Genes Undergo Diversity-Enhancing Evolution in a Fungal Species Complex. Genome Biology and Evolution 18, doi:10.1093/gbe/evag185 (2026).

38 Torres, A. M., Weeden, N. F. & Martín, A. Linkage among isozyme, RFLP and RAPD markers in Vicia faba. Theoretical and Applied Genetics 85, 937–945, doi:10.1007/BF00215032 (1993).

39 Di Pietro, A. et al. Endopolygalacturonase PG1 in Different Formae Speciales of Fusarium oxysporum. Applied and Environmental Microbiology 64, 1967–1971, doi:10.1128/AEM.64.5.1967-1971.1998 (1998).

40 Joska, T. M., Mashruwala, A., Boyd, J. M. & Belden, W. J. A universal cloning method based on yeast homologous recombination that is simple, efficient, and versatile. Journal of Microbiological Methods 100, 46–51, 10.1016/j.mimet.2013.11.013 (2014).

41 Redkar, A. et al. Conserved secreted effectors determine endophytic growth and multi-host plant compatibility in a vascular wilt fungus. The Plant Cell 34, 3214–3232, doi: 10.1093/plcell/koac174 (2022).

42 Punt, P. J., Oliver, R. P., Dingemanse, M. A., Pouwels, P. H. & van den Hondel, C. A. M. J. J. Transformation of Aspergillus based on the hygromycin B resistance marker from Escherichia coli. Gene 56, 117–124, 10.1016/0378-1119(87)90164-8 (1987).

43 Nordzieke, D. E., Fernandes, T. R., El Ghalid, M., Turra, D. & Di Pietro, A. NADPH oxidase regulates chemotropic growth of the fungal pathogen Fusarium oxysporum towards the host plant. New Phytol 224, 1600–1612, doi:10.1111/nph.16085 (2019).

44 Lander, R. J., Winters, M. A., Meacle, F. J., Buckland, B. C. & Lee, A. L. Fractional precipitation of plasmid DNA from lysate by CTAB. Biotechnology and Bioengineering 79, 776–784, 10.1002/bit.10335 (2002).

45 Silva, D., Santos, G., Barroca, M. & Collins, T. in PCR: Methods and Protocols (ed Lucília Domingues) 87–100 (Springer New York, 2017).

46 López-Berges, M. S., Rispail, N., Prados-Rosales, R. C. & Di Pietro, A. A Nitrogen Response Pathway Regulates Virulence Functions in Fusarium oxysporum via the Protein Kinase TOR and the bZIP Protein MeaB The Plant Cell 22, 2459–2475, doi:10.1105/tpc.110.075937 (2010).

47 Mattern, I., Punt, P. & Van den Hondel, C. A vector of Aspergillus transformation conferring phleomycin resistance. Fungal Genet Newsl 35, 25 (1988).

48 Corral-Ramos, C., Roca, M. G., Di Pietro, A., Roncero, M. I. G. & Ruiz-Roldán, C. Autophagy contributes to regulation of nuclear dynamics during vegetative growth and hyphal fusion in Fusarium oxysporum. Autophagy 11, 131–144, doi:10.4161/15548627.2014.994413 (2015).

49 Berensmeier, S. Magnetic particles for the separation and purification of nucleic acids. Applied Microbiology and Biotechnology 73, 495–504, doi:10.1007/s00253-006-0675-0 (2006).

50 Pfafl, M. W. A new mathematical model for relative quantification in real-time RT–PCR. Nucleic Acids Research 29, e45–e45, doi:10.1093/nar/29.9.e45 (2001).

51 Palos-Fernández, R. et al. Copper acquisition is essential for plant colonization and virulence in a root-infecting vascular wilt fungus. PLOS Pathogens 20, e1012671, doi:10.1371/journal.ppat.1012671 (2024).

