## Supplementary Materials and Methods for "Killer meiotic driver prevents chromosome loss and enhances fitness in a fungal pathogen"

#### Whole-genome synteny between MRL8996 and Fol4287

The chromosome-scale synteny between MRL8996 (GCA_009746015.2) and Fol4287 (GCA_000149955.2) was analysed using minimap2[1] version 2.26 in assembly-to-assembly mode (`-x asm20 -t 8`). Alignment blocks shorter than 5,000 base pairs were excluded, and the remaining blocks were aggregated by chromosome pair based on the sum of aligned block lengths. This method allowed for the determination of the dominant correspondence for each MRL8996 chromosome based on total aligned sequences rather than any single alignment.

Two reference organisations were utilised. The first used the fifteen RefSeq chromosome accessions of Fol4287 (NC_030986.1–NC_031000.1) mapped to their respective chromosome numbers. The final figure was based on the reorganised JGI Fol4287 assembly, where the arcs are arranged with core chromosomes first, followed by accessory chromosomes. In this representation, the region between chromosomes 3 and 6 is depicted by two separate contigs, labelled as chromosome 3 and accessory arc S2. MRL8996 chromosomes were named in descending order of size and plotted in reverse order to ensure that syntenic partners face each other across the circle.

Circular synteny plots were created in R using the `circlize`[2] package. Each arc of MRL8996 was coloured according to its principal partner in Fol4287, with ribbons coloured by their respective Fol4287 chromosome-of-origin at 40% opacity. The accessory chromosomes of both genomes were displayed in grey to distinguish the conserved core from the lineage-specific compartments. The corresponding pairwise coverage matrix (Supplementary Table S5) details the percentage of the query chromosome covered by alignments to each reference chromosome.

Chromosome correspondence between the two genomes was determined independently from the NUCmer alignments used for interval extraction by aggregating filtered alignment blocks for each chromosome pair (Supplementary Table S6). Eleven chromosomes of MRL8996 correspond one-to-one with a core chromosome of Fol4287, showing 76–89% coverage of the reference chromosome. In contrast, the accessory chromosomes of Fol4287 (Chr03, Chr06, Chr14, Chr15) do not have counterparts in MRL8996, aligning to less than 8% of their length.

**Whole-genome alignment and extraction of chromosome 12 and chromosome 10 intervals**

A total of 1,252 publicly available genomic assemblies from the *Fusarium* species complex were downloaded from NCBI GenBank and RefSeq. This included the chromosome-level assembly of *F. oxysporum* strain MRL8996[3]. Throughout this study, the assembly of *F. oxysporum* f. sp. *lycopersici* strain 4287 (GCF_000149955.1, ASM14995v2) served as the reference genome. Specifically, we focused on Fol4287 chromosome 12 of this strain (RefSeq NC_030997.1, GenBank CM000600.1; 2,233,407 bp), which contains the *Starship* element and the *spok* homolog examined in our research. For comparative analysis, Fol4287 chromosome 10 (NC_030995.1; 2,896,840 bp) was used as a core chromosome control of similar size. The forma specialis and species assignments for each assembly were obtained from the NCBI BioSample and assembly metadata (Supplementary Table S7).

Each assembly was aligned against the Fol4287 reference genome using NUCmer[4] (MUMmer 3.1) with default seed and clustering parameters. Raw delta alignments were filtered using the command `delta-filter -i 85 -l 5000 -q`, which retained alignment blocks with at least 85% nucleotide identity and a minimum length of 5,000 bp. For each query position, only the best reference match was kept. Filtered alignments were converted to tabular coordinates using `show-coords -r -T -H`.

For each target chromosome, alignment blocks anchored to the corresponding reference accession were extracted from the coordinate tables, sorted by query contig and start position, and merged using BEDTools[5] (v2.31.1) with the command `merge -d 10000`. This process joined syntenic blocks that were separated by less than 10 kb into a single interval. The merged intervals were retrieved from the query assembly with `bedtools getfasta`, resulting in one multi-FASTA file of chromosome-12 syntenic sequences and one of chromosome-10 syntenic sequences for each genome. Chromosome 10 was processed using the same method on the same set of filtered delta files to ensure that identity and length thresholds were strictly comparable between the two chromosomes.

For every assembly, this procedure recorded the number of contributing query contigs, the total extracted length, the length-weighted mean nucleotide identity, and the fraction of the reference chromosome recovered (Supplementary Table S8). Assemblies recovering less than 70% of the reference chromosome were excluded from all downstream phylogenetic analyses; 421 assemblies passed this filter for chromosome 12, while 468 passed for chromosome 10.

**Gene prediction, orthology inference and phylogenomic reconstruction**

The syntenic intervals of chromosome 12 and chromosome 10 from each genome that passed the 70% recovery filter were annotated ab initio using AUGUSTUS[6] v3.5.0 with the options `--species=fusarium --gff3=on`. From the resulting GFF3 files, we extracted the predicted protein sets, yielding 421 proteomes for Chromosome 12 and 468 proteomes for chromosome 10.

Orthogroups were inferred using OrthoFinder[7] v2.5.5 in multiple-sequence alignment mode (`-M msa`), which utilises DIAMOND for all-versus-all similarity searches and MAFFT for orthogroup alignment. The concatenated single-copy orthogroup alignments generated by OrthoFinder were used to create phylogenomic supermatrices.

Maximum-likelihood trees were constructed with IQ-TREE[8] under the LG+I+G4 model, using 1,000 ultrafast bootstrap replicates (`-m LG+I+G4 -B 1000 -T 16`). The trees were visualised as circular (fan) layouts and midpoint-rooted in R v4.3.3 using the `ggtree`[9] and `ggtreeExtra`[10] packages. These trees were annotated with concentric rings representing forma specialis, chromosome recovery class, presence in the reciprocal tree, and mean nucleotide identity of the extracted chromosome. In the Figure for chromosome 10, the colour scale for nucleotide identity is set to a minimum of approximately 97.4% instead of zero, as the identity range for this core chromosome is too narrow to be visually distinguished otherwise.

### Detection of the *Starship* element, its Captain, and Spok

For homology searches, we utilized three reference query sequences: (i) the *Starship* element of Fol4287, defined by the interval NC_030997.1:2,051,479–2,144,131 (92,653 bp); (ii) the associated Captain tyrosine recombinase, FOXG_14754 (NCBI protein XP_018254392.1; 829 amino acids); and (iii) the Fol4287 Spok homolog, FOXG_14774 (XP_018254443.1; 727 amino acids).

We converted the extracted chromosome 12 intervals of each genome into a BLAST nucleotide database (one database per genome) using `makeblastdb` (BLAST+ v2.12.0[11]). We conducted three searches in a common coordinate space: the *Starship* element was searched using `blastn` ($E \leq 1 \times 10^{-20}$, `-max_target_seqs 50`), while both the Captain and the Spok protein were searched using `tblastn` ($E \leq 1 \times 10^{-10}$, `-max_target_seqs 20`).

For each query, we merged all high-scoring segment pairs in query coordinates before computing coverage, ensuring that multi-block and multi-exon hits were not double-counted. The sequence identity was reported as the alignment-length-weighted mean over the contributing blocks (Supplementary Tables S8 and S9)

The presence of each feature was determined by evaluating both merged query coverage and sequence identity. Specifically, an 'element' was considered present if it exhibited a merged query coverage of at least 50% and at least 80% nucleotide identity (Supplementary Table S10). For 'Captain' and 'Spok', presence was assessed at the protein level: Captain required a merged coverage of at least 70% with a minimum of 35% amino acid identity, while Spok demanded a higher standard, requiring at least 80% merged coverage and a minimum of 70% amino acid identity (Supplementary Tables S11 and S12).

A Spok hit was classified as `INSIDE_STARSHIP` if its subject interval was within the merged element footprint, which is extended by a 2,000 bp pad on either side. It was classified as `ORPHAN_ON_CHR12` if it met the presence thresholds but lay outside that footprint. The completeness of the element was summarised into four descriptive categories: intact (≥ 75% coverage and ≥ 75% identity), partial (50–74% coverage), fragment/relic (> 0 but < 50% coverage), and absent (Supplementary Tables S12 and S13).

The status of Captain was categorised into three classes based on whether a qualifying hit was present and located within 5 kb of the element start, present but internal or misaligned, or fragmented/absent. Additionally, any genomes with less than 40% recovery of chromosome 12 were flagged in the summary to prevent confusion between true absence and poor recovery of the chromosome itself.

**Functional annotation and Gene Ontology enrichment in MRL8996**

Gene models for MRL8996 were obtained from the polished annotation of the assembly. Gene Ontology terms were sourced from the JGI gene catalogue's GO assignment table for this strain, which included 31,036 protein-to-term assignments. Genes were allocated to chromosomes using the scaffold-to-chromosome mapping of the assembly. This assignment was independently verified in this study by aligning the polished scaffolds against Fol4287 using minimap2[1] (`-x asm20`, MAPQ ≥ 10, blocks ≥ 5 kb). The alignment confirmed that the dominant reference chromosome for each mapped scaffold was consistent with the declared assignments, as indicated in Supplementary Table S14, with 11 out of 11 assigned scaffolds showing concordance.

Enrichment was tested with a one-sided Fisher exact test contrasting, for each GO term, the number of annotated genes carrying that term on the chromosome of interest against the number on all remaining annotated chromosomes. Terms supported by fewer than two genes on the target chromosome were not tested.

We executed two complementary analyses to ensure a comprehensive assessment of the data. The first was focused on chromosome 10. For this analysis, we specifically examined the GO terms associated with MRL8996 chromosome 10, which is syntenic to Fol4287 chromosome 12. We then compared these terms to those in the rest of the annotated genome. To account for the potential of false discoveries, we applied the Benjamini–Hochberg procedure for multiple testing corrections. Only terms with a q-value of less than 0.05 were reported, culminating in the identification of 19 significant terms (Supplementary Table S14). The second analysis took a broader approach, encompassing a genome-wide profile. Here, we applied the same Fisher exact test across all eleven assigned chromosomes, looking for terms that exhibited both a p-value of less than 0.01 and a fold enrichment of at least 1.5. This analysis yielded 38 terms across five chromosomes, with corresponding Benjamini–Hochberg q-values provided for reference (Supplementary Table S15).

GO term names and ontology aspects (biological process, molecular function, cellular component) were retrieved programmatically from the EBI QuickGO[12] service.

**Structural comparison of the SPOK proteins**

Previously, Vogan et al., 2021[13] identified the catalytic lysine K240 of the PD-(D/E)xK nuclease domain of *Podospora anserina* Spok3 as essential for the spore-killing phenotype; mutating K240 to K240A abolished this phenotype. To investigate whether this residue and its structural context are conserved in *Fusarium*, the *F. oxysporum* Spok homolog FOXG_14774 (XP_018254443.1; 727 aa) and *P. anserina* Spok3 (UniProt A0A516F180; 745 aa) were modelled and compared.

Both proteins were locally folded with ColabFold[14] with identical settings (`--model-type alphafold2_ptm --msa-mode mmseqs2_uniref_env --num-models 5 --num-recycle 3 --rank plddt --num-relax 0`), and the top-ranked model of each was retained. The mean pLDDT scores were 77.3 for the *F. oxysporum* protein homolog and 74.0 for SPOK3. As an independent verification of the Spok3 model, the ColabFold model was compared to the AlphaFold Database[15] entry for A0A516F180, which has a mean pLDDT of 75.8.

The *F. oxysporum* Spok model was superimposed onto the locally folded SPOK3 model using ChimeraX[16] MatchMaker under default sequence-plus-secondary-structure settings. Catalytic and DNA-binding residues were mapped between the two proteins through the pairwise sequence alignment underlying the superposition.

**DNA extraction and Nanopore library preparation and sequencing**

For genomic DNA isolation, mycelium from MRL8996 and MRL8996 -chr10 was collected by filtration through a Monodour nylon filter and freeze-dried overnight. High-molecular-weight (HMW) DNA was extracted following the protocol of Chavarro-Carrero et al. (2021)[17]. DNA quality, size, and quantity were assessed using NanoDrop, Qubit, and gel electrophoresis. Library preparation with the Native Barcoding Ligation Sequencing Kit (SQK-NBD114.24) was performed with ~1.5 µg HMW DNA according to the manufacturer's instructions (Oxford Nanopore Technologies, Oxford, UK). An R10.4.1 flow cell (Oxford Nanopore Technologies, Oxford, UK) was loaded and run for 24 h, and subsequent basecalling was performed using Dorado (version 7.11.2; Oxford Nanopore Technologies, Oxford, UK).

**Copy-number analysis of the MRL8996 chromosome-loss mutant**

Copy-number variation was assessed by whole-genome Oxford Nanopore resequencing of two strains of *F. oxysporum* MRL8996: the wild-type isolate and a derivative that had lost the chromosome homologous to chromosome 12 of Fol4287, designated MRL8996 −chr10 throughout.

Reads of each sample were aligned to the reference with minimap2[1] in Nanopore mode (-ax map-ont), and the resulting SAM streams were converted, coordinate-sorted and indexed with samtools. Per-base sequencing depth was extracted with samtools depth -a, so that positions with zero coverage are reported explicitly, and every chromosome contributes a complete positional record; this is required for the windowing step, in which absent positions must be distinguished from missing lines.

Depth records were reduced to non-overlapping 10-kb windows tiling each chromosome, each window being summarised by the median of its per-base depths; the median was preferred over the mean because it is insensitive to the short, very high-depth spikes produced by collapsed repeats. Windows containing no depth record were assigned a value of zero.

Normalisation was conducted in three sequential steps. First, to account for differences in sequencing library yield, absolute read depth was converted to relative coverage. This was done by dividing each sample's window vector by the median of its own non-zero values. This per-sample scaling ensures that the mode of the single-copy region for each sample is standardised to a value of 1.

Second, a wild-type copy-number correction was applied to prevent regions with natural duplications in the wild type from being misinterpreted as spurious gains in the mutant. The scaled wild-type vector was rounded to the nearest integer and floored at 1 to establish a per-window copy-number baseline. Both samples were then divided by this baseline. As a result, any window that maintained its wild-type copy number took on a value of 1 in both samples, regardless of the absolute copy number.

Finally, for windows located on core chromosomes, this corrected per-sample value was retained. However, windows on accessory chromosomes were further divided by the corrected wild-type value to represent them as a mutant-to-wild-type ratio. Any infinite or undefined values resulting from division by zero-coverage windows were adjusted to 1.

To visualise these data, two genome-wide coverage tracks were created as bar charts plotted along the concatenated window index. Chromosome boundaries were explicitly marked, and a baseline reference was set at unity. These visualisations included a profile track displaying the corrected per-sample coverage for each strain, and a ratio track representing the coverage of each strain relative to the corrected wild type. Both tracks were scaled to a common range of 0–5× to facilitate direct comparison of whole-chromosome losses and gains across different panels (Supplementary Tables S16 and S17).

**REFERENCES**

1. Li, H. Minimap2: Pairwise alignment for nucleotide sequences. *Bioinformatics* **34**, (2018).

2. Gu, Z., Gu, L., Eils, R., Schlesner, M. & Brors, B. Circlize implements and enhances circular visualization in R. *Bioinformatics* **30**, (2014).

3. Doddi, A., Puebla Planas, G., López-Berges, M. S. & Di Pietro, A. A near-complete genome assembly of the *Fusarium oxysporum* keratitis isolate MRL8996. Preprint at https://doi.org/10.64898/2026.07.06.736765 (2026).

4. Kurtz, S. *et al.* Versatile and open software for comparing large genomes. *Genome Biol.* **5**, (2004).

5. Quinlan, A. R. & Hall, I. M. BEDTools: A flexible suite of utilities for comparing genomic features. *Bioinformatics* **26**, (2010).

6. Stanke, M. *et al.* AUGUSTUS: A b initio prediction of alternative transcripts. *Nucleic Acids Res.* **34**, (2006).

7. Emms, D. M. & Kelly, S. OrthoFinder: Phylogenetic orthology inference for comparative genomics. *Genome Biol.* **20**, (2019).

8. Nguyen, L. T., Schmidt, H. A., Von Haeseler, A. & Minh, B. Q. IQ-TREE: A fast and effective stochastic algorithm for estimating maximum-likelihood phylogenies. *Mol. Biol. Evol.* **32**, (2015).

9. Yu, G., Smith, D. K., Zhu, H., Guan, Y. & Lam, T. T. Y. ggtree: an r package for visualization and annotation of phylogenetic trees with their covariates and other associated data. *Methods Ecol. Evol.* **8**, (2017).

10. Attali, D. & Baker, C. ggExtra: Add Marginal Histograms to 'ggplot2', and More 'ggplot2' Enhancements. *R J.* (2022).

11. Camacho, C. *et al.* BLAST+: Architecture and applications. *BMC Bioinformatics* **10**, (2009).

12. Binns, D. *et al.* QuickGO: A web-based tool for Gene Ontology searching. *Bioinformatics* **25**, (2009).

13. Vogan, A. A. *et al.* The Enterprise, a massive transposon carrying Spok meiotic drive genes. *Genome Res.* **31**, (2021).

14. Mirdita, M. *et al.* ColabFold: making protein folding accessible to all. *Nat. Methods* **19**, (2022).

15. Varadi, M. *et al.* AlphaFold Protein Structure Database in 2024: providing structure coverage for over 214 million protein sequences. *Nucleic Acids Res.* **52**, (2024).

16. Meng, E. C. *et al.* UCSF ChimeraX: Tools for structure building and analysis. *Protein Science* **32**, (2023).

17. Chavarro-Carrero, E. A. *et al.* Comparative genomics reveals the in planta-secreted Verticillium dahliae Av2 effector protein recognized in tomato plants that carry the V2 resistance locus. *Environ. Microbiol.* **23**, (2021).
